# Combinatorial control of biofilm development by quorum-sensing and nutrient-sensing regulators in *Pseudomonas aeruginosa*

**DOI:** 10.1101/2022.09.27.509822

**Authors:** Gong Chen, Georgia Fanouraki, Aathmaja Anandhi Rangarajan, Bradford T. Winkelman, Jared T. Winkelman, Christopher M. Waters, Sampriti Mukherjee

## Abstract

The human pathogen *Pseudomonas aeruginosa*, a leading cause of hospital-acquired infections, inhabits and forms sessile antibiotic-resistant communities called biofilms in a wide range of biotic and abiotic environments. In this study, we examined how two global sensory signaling pathways – the RhlR quorum-sensing system and the CbrA/CbrB nutritional adaptation system – intersect to control biofilm development. Previous work has shown that individually these two systems repress biofilm formation. Here, we used biofilm analyses, RNA-seq, and reporter assays to explore the combined effect of information flow through RhlR and CbrA on biofilm development. We find that the Δ*rhlR*Δ*cbrA* double mutant exhibits a biofilm morphology and an associated transcriptional response distinct from wildtype and the parent Δ*rhlR and* Δ*cbrA* mutants indicating codominance of each signaling pathway. The Δ*rhlR*Δ*cbrA* mutant rapidly gains suppressor mutations that map to the carbon catabolite repression protein Crc. The combined absence of RhlR and CbrA leads to drastic reduction in the abundance of the Crc antagonist small RNA CrcZ. Thus, CrcZ acts as the molecular convergence point for quorum- and nutrient-sensing cues. Furthermore, in the absence of antagonism by CrcZ, Crc promotes the expression of biofilm matrix components – Pel exopolysaccharide, and CupB and CupC fimbriae. Therefore, this study uncovers a regulatory link between nutritional adaption and quorum sensing with potential implications for anti-biofilm targeting strategies.

**AUTHOR SUMMARY:** Bacterial pathogens often form multicellular communities encased in an extra cytoplasmic matrix called biofilms as a virulence strategy. Biofilm development is controlled by various environmental stimuli that are decoded and converted into appropriate cellular responses. How information from two or more stimuli is integrated is poorly understood. Using *Pseudomonas aeruginosa* biofilm formation as a model, we studied the intersection of two global sensory signaling pathways – quorum sensing and nutritional adaptation. We find parallel regulation by each pathway that converges on the abundance of a small RNA. Thus, we describe a regulatory link between *P. aeruginosa* quorum-sensing and nutritional adaptation pathways that allows integration of information from each system into the control of biofilm development. These results expand our understanding of the genetic regulatory strategies that allow *P. aeruginosa* to successfully colonize host during chronic infections.

## INTRODUCTION

Bacteria predominantly exist in structured communities called biofilms. Biofilms are defined as aggregates of cells that are embedded in a matrix made of extracellular polymeric substances (EPS) including exopolysaccharides, proteins, lipids and nucleic acids (Hall-Stoodley et al. 2004; Flemming et al. 2016; Karygianni et al. 2020). The EPS is crucial for the emergent properties of biofilms such as superior resilience to environmental stresses like antimicrobials and host immune responses. Biofilm formation is a dynamic process that is governed by various intracellular and exogenous stimuli. Often, two-component signaling (TCS) systems integrate and relay the information contained in sensory cues into the control of biofilm formation (Liu et al., 2018; Prüß, 2017). TCSs are typically composed of sensor histidine kinases (HK) and partner response regulators (RR) that, via phosphorylation cascades, couple stimulus sensing to appropriate changes in behavior (Capra and Laub, 2012; Liu et al., 2019).

The opportunistic pathogen *Pseudomonas aeruginosa* forms biofilms in diverse free-living environments, such as soil and water and in host-associated environments, such as burn wounds, lungs of cystic fibrosis (CF) patients and plant tissues (Pendleton et al. 2013; Markou and Apidianakis, 2014; Manfredi et al. 2000; Kim et al. 2015). Accordingly, *P. aeruginosa* encodes a large suite of >60 TCS systems that allow it to respond to diverse external cues and adapt to a wide range of infection sites and environmental conditions (Wang et al. 2021). One such TCS is the CbrA/CbrB TCS that is involved in nutritional adaptation and was first described in *P. aeruginosa* as a regulatory system involved in the hierarchical utilization of various carbon sources (Monteagudo-Cascales et al. 2022; Sonnleitner et al. 2009; Yeung et al. 2011; Sonnleitner et al. 2018). CbrA represents a non-canonical sensor HK as it can function as a histidine transporter via its large N-terminal transmembrane region called the SLC5 (SoLute Carrier 5) domain as well as a HK via its C-terminal catalytic core consisting of a DHp (Dimerization and Histidine Phosphotransfer) domain and a catalytic ATP-Binding domain (Monteagudo-Cascales et al. 2022). CbrA has two putative sensor domains – SLC5 and PAS, yet the sensory stimulus activating its kinase is unknown. CbrA kinase phosphorylates its cognate RR CbrB (Fig. 1A).

**Fig. 1:**
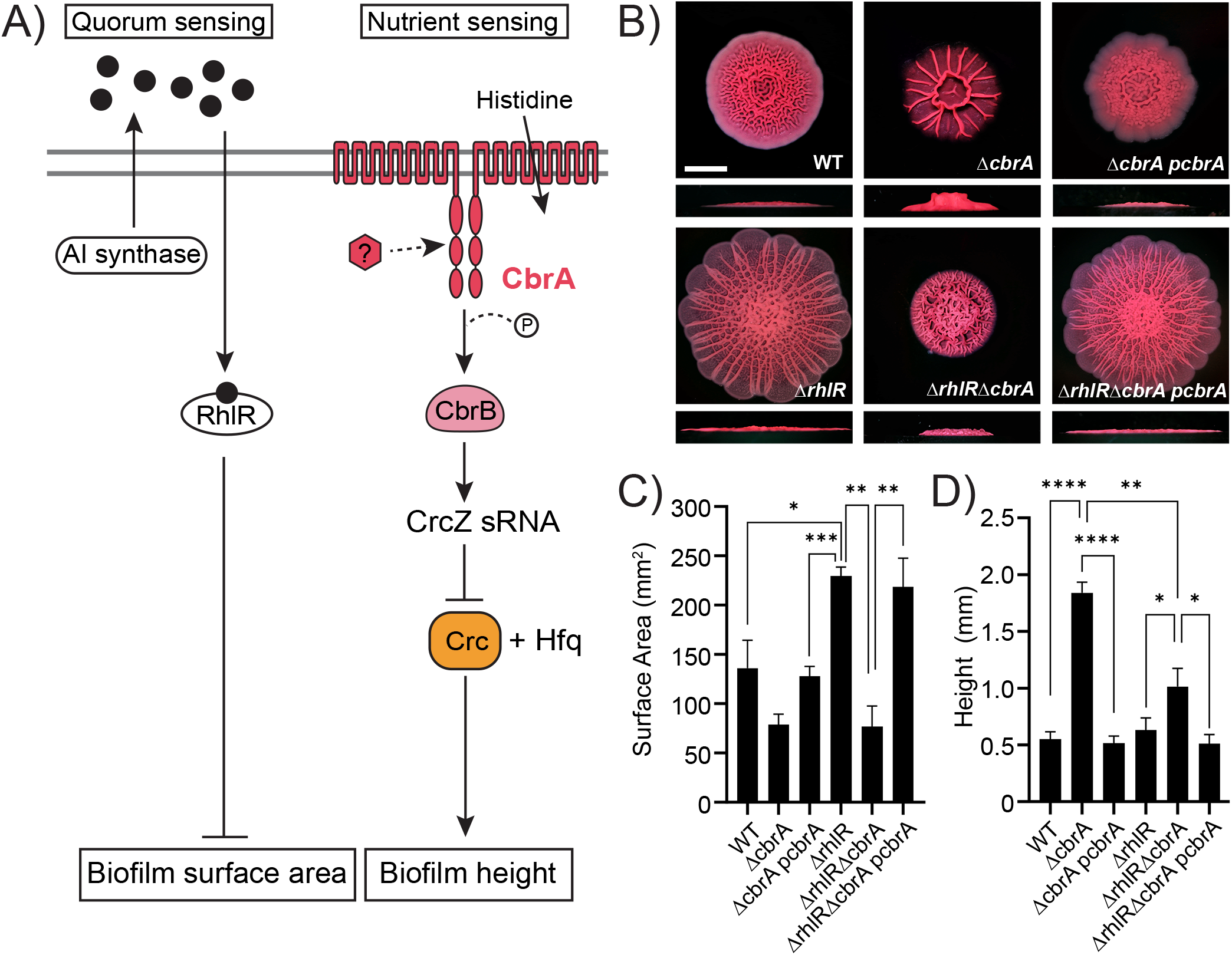
*P. aeruginosa* Δ*cbrA* and Δ*rhlR* mutants have distinct hyper-rugose biofilm phenotypes. A) Schematic of the RhlR quorum sensing and CbrA nutrient sensing pathways. The two gray horizontal lines represent the cytoplasmic membrane, black circles represent autoinducer sensed by RhlR, red hexagon represents the unknown signal that activates CbrA sensor kinase. CbrA also functions as a transporter of histidine but its kinase function appears to be independent of histidine transport (Monteagudo-Cascales et al. 2022). B) Colony biofilm phenotypes of WT PA14 and the designated mutants on Congo red agar medium after 120 h of growth. Scale bar, 5 mm. C) Colony biofilm surface area quantitation for the indicated strains after 120 h of growth. Error bars represent standard deviation of three independent experiments. Statistical significance was determined using two-tailed *t*-test with Welch’s correction in GraphPad Prism software. *** P <0.001, ** P <0.01, * P <0.05. D) Colony biofilm height quantitation for the indicated strains after 120 h of growth. Error bars represent standard deviation of three independent experiments. Statistical significance was determined using two-tailed *t*-test with Welch’s correction in GraphPad Prism software. **** P <0.0001, ** P <0.01, * P <0.05.

CbrB is an NtrC class transcription factor that possesses a REC receiver domain, a AAA+ ATPase domain and a HTH (Helix-Turn-Helix) DNA-binding domain and activates transcription from σ^54^ dependent promoters. The genes *cbrA* and *cbrB* are encoded in the same transcriptional unit. CbrB activates the expression of the small RNA CrcZ that is encoded immediately downstream of *cbrB* in the genome (Kambara et al., 2018, Sonnleitner E, et al. 2018; Fig. 1A). CrcZ small RNA sequesters complexes of the catabolite repression protein Crc with Hfq to antagonize Crc function in the carbon catabolite repression (CCR) process termed “reverse diauxie” or reverse CCR (rCCR, Park et al., 2020, McGill et al., 2021). In the presence of preferred carbon sources, CrcZ mediated antagonism is relieved and Crc/Hfq post-transcriptionally inhibit the expression of enzymes involved in catabolism of non-preferred carbon sources. In addition to the metabolic regulation of carbon and nitrogen utilization in *P. aeruginosa*, Cbr TCS plays an important role in various virulence-associated processes including biofilm formation and antibiotic resistance (Yeung, et al. 2011).

Another sensory cue commonly detected by bacteria is their population density via the chemical communication process called quorum sensing (Rutherford and Bassler, 2016). Quorum sensing relies on production of extracellular signaling molecules called autoinducers and their subsequent detection by cognate receptors (Miller and Bassler, 2001; Mukherjee and Bassler, 2019), and thereby allows groups of bacteria to coordinate their gene expression patterns in response to changes in population density to exhibit collective behaviors. Quorum sensing is crucial for biofilm development and virulence in *P. aeruginosa* (Davies et al., 1998; Rumbaugh et al., 2000; Mukherjee et al., 2017). The *P. aeruginosa* quorum-sensing circuit consists of two canonical LuxI/R pairs: LasI/R and RhlI/R (Seed et al. 1995; Pearson et al. 1995; Brint and Ohman,1995). LasI produces and LasR responds to the autoinducer 3OC12-homoserine lactone (3OC12-HSL) while RhlR binds to the autoinducer C4-HSL, the product of RhlI. *P. aeruginosa* LasI/R promotes biofilm formation (Davies *et al*, 1998; Sakuragi & Kolter, 2007). Conversely, we recently discovered that the *P. aeruginosa* quorum-sensing receptor RhlR represses biofilm formation (Mukherjee et al. 2017; Fig. 1A).

Bacteria encounter multiple sensory cues simultaneously and must integrate information from each cue into a response. In this study, we examine how the CbrA/CbrB signaling pathway intersects with the RhlR dependent quorum sensing system to control biofilm development. We find that the absence of both signals – quorum sensing via RhlR and nutrient sensing via CbrA results in (a) a biofilm morphology and an associated transcriptional response that indicates codominance of each signaling pathway, and (b) significant reduction in the levels of the small RNA CrcZ. We further find that the formation of a hyper-rugose biofilm in the Δ*rhlR*Δ*cbrA* mutant appears to be unstable and readily gives rise to spontaneous suppressor mutations in the *crc* gene encoding for the carbon catabolite repression protein Crc. Transcriptomic profiling of wildtype and mutant *P*. *aeruginosa* biofilms and reporter assays demonstrate that Crc promotes the expression of Pel exopolysaccharide and Cup fimbriae components of the biofilm matrix. Thus, our work uncovers a new role for Crc as a master regulator of biofilm formation that allows the integration of quorum and nutritional cues.

## RESULTS

### CbrA-mediated nutrient-sensing and RhlR-mediated quorum-sensing co-dominantly control biofilm development

To study the role of the CbrA/CbrB TCS in *P*. *aeruginosa* biofilm formation, we first generated in-frame marker-less deletion of the *cbrA* gene in the wildtype (WT) *P*. *aeruginosa* UCBPP-PA14 (hereafter called PA14) background. PA14 exhibits a characteristic rugose-center/smooth-periphery colony biofilm phenotype on Congo red biofilm medium while the Δ*cbrA* mutant exhibits a distinct colony biofilm phenotype with decreased surface area coverage and increased height, defined as the maximum vertical rise from the base of the colony, when compared to WT (Fig. 1B-D, S1). CbrA is a non-canonical membrane-bound sensor kinase that can also function as a histidine transporter (Zhang et al., 2015; Monteagudo-Cascales et al., 2019; Wirtz et al. 2020). Therefore, to uncouple its two functions and determine whether the hyper-rugose biofilm phenotype of the Δ*cbrA* mutant is a consequence of the absence of information flow from CbrA kinase to CbrB response regulator, we generated a Δ*cbrB* mutant. The Δ*cbrB* mutant produces hyper-rugose biofilms identical to the Δ*cbrA* mutant (Fig. S2). In addition, we built a *cbrA^H766A^* mutant where the histidine residue that undergoes autophosphorylation is changed to alanine to abolish kinase function of CbrA. Similar to the Δ*cbrB* strain, the *cbrA^H766A^* mutant has a biofilm phenotype that resembles the Δ*cbrA* mutant (Fig. S2). Introduction of a plasmid expressing *cbrA* under its native promoter to the Δ*cbrA* and *cbrA^H766A^* mutants restored biofilm formation to wildtype (WT) levels (Fig. 1B-D, S2). These analyses indicate that CbrA kinase activity is required for repression of biofilm development in *P. aeruginosa*.

Production of the extracellular matrix is a defining feature of biofilms, and the matrix composition can vary depending on the bacterial species. *P. aeruginosa* biofilm matrix has been reported to be composed of three different exopolysaccharides – Pel, Psl and alginate – depending on the strain (reviewed in Jiang et al., 2021). In PA14, Pel is the major exopolysaccharide as Psl is not made by this strain and alginate is not a significant component of the biofilm matrix under commonly used laboratory growth conditions (Jennings et al. 2015). Pel biosynthetic enzymes are encoded by the *pelABCDEFG* operon. Here, we assessed the contribution of Pel to the Δ*cbrA* mutant biofilms by comparing the Δ*pelA* and Δ*cbrA*Δ*pelA* double mutants. Absence of PelA abolishes formation of colony biofilms and pellicles (Fig. S3 and S4). Both the Δ*pelA* and Δ*cbrA*Δ*pelA* mutants had completely smooth colony morphologies and lower biofilm height compared to WT (Fig. S4A-C). Furthermore, in a solid surface-associated (SSA) biofilms assay, they showed significantly less crystal violet staining compared to the Δ*cbrA* mutant (Fig. S4D). Thus, the Pel exopolysaccharide is required for the Δ*cbrA* mutant to exhibit its characteristic hyper-rugose biofilm phenotype.

We have previously reported that the *P. aeruginosa* quorum-sensing receptor RhlR represses colony biofilm development and the hyper-rugosity conferred by the absence of RhlR requires the primary biofilm matrix exopolysaccharide Pel (Fig. 1, Mukherjee et al., 2017). Intriguingly, although both Δ*rhlR* and Δ*cbrA* mutants appear to be hyper-rugose compared to WT PA14, the Δ*cbrA* mutant exhibits increased biofilm height as opposed to the increased surface area coverage of the Δ*rhlR* mutant (Fig. 1B-D, S1). To explore the combined effect of Rhl and Cbr pathways on biofilm formation, we deleted *rhlR* and *cbrA* together. The Δ*rhlR*Δ*cbrA* double mutant formed colony biofilms that were markedly distinct from those of the single Δ*rhlR* and Δ*cbrA* mutants, i.e., had significantly lower surface coverage than Δ*rhlR* mutant and lower height than Δ*cbrA* mutant (Fig. 1B-D, S1). Introduction of a plasmid expressing *cbrA* under its native promoter to the Δ*rhlR*Δ*cbrA* double mutant resulted in a hyper-rugose biofilm that resembles the Δ*rhlR* mutant in terms of biofilm morphology, surface area and height (Fig. 1B-D). We conclude that RhlR and CbrA control different properties, surface area coverage and vertical rise, respectively, of a growing biofilm.

### Loss-of-function mutations in *crc* affects biofilm development

The Δ*rhlR*Δ*cbrA* double mutant formed unstable biofilms that readily gave rise to suppressor flares that allowed spreading of the biofilm similar to the Δ*rhlR* mutant (Fig. 2A). We isolated 12 spontaneously arising suppressor mutants from Δ*rhlR*Δ*cbrA* colony biofilms and analyzed them by whole genome sequencing. Ten suppressors contained deletions, insertions or missense mutations in the *crc* gene, while the remaining two suppressors harbored mutations in the *rplP* and *rplC* genes that encode for 50S ribosomal protein L16 and 50S ribosomal protein L3, respectively (Fig. 2A, Table S1). We generated Δ*crc* single, Δ*rhlR*Δ*crc* and Δ*cbrA*Δ*crc* double, and Δ*rhlR*Δ*cbrA*Δ*crc* mutants and analyzed their biofilm forming behaviors. Absence of Crc showed little effect in WT and Δ*rhlR* backgrounds but substantially altered the biofilms of the Δ*cbrA* and Δ*rhlR*Δ*cbrA* mutants (Fig. 2B-D). Specifically, deletion of *crc* abolished the vertical rise of the Δ*cbrA* mutant colony biofilm (Fig. 2B-D) and led to increased surface area coverage of Δ*rhlR*Δ*cbrA* double mutant colony biofilms (Fig. 2B-D, S1). Introduction of a plasmid expressing *crc* under its native promoter to the Δ*rhlR*Δ*cbrA*Δ*crc* triple mutant resulted in colony biofilm phenotype identical to the Δ*rhlR*Δ*cbrA* double mutant validating our deletion analyses (Fig. 2B-D). We conclude that Crc promotes colony biofilm height when CbrA is inactive and restricts colony biofilm spreading when signal transduction via both RhlR and CbrA is absent.

**Fig. 2:**
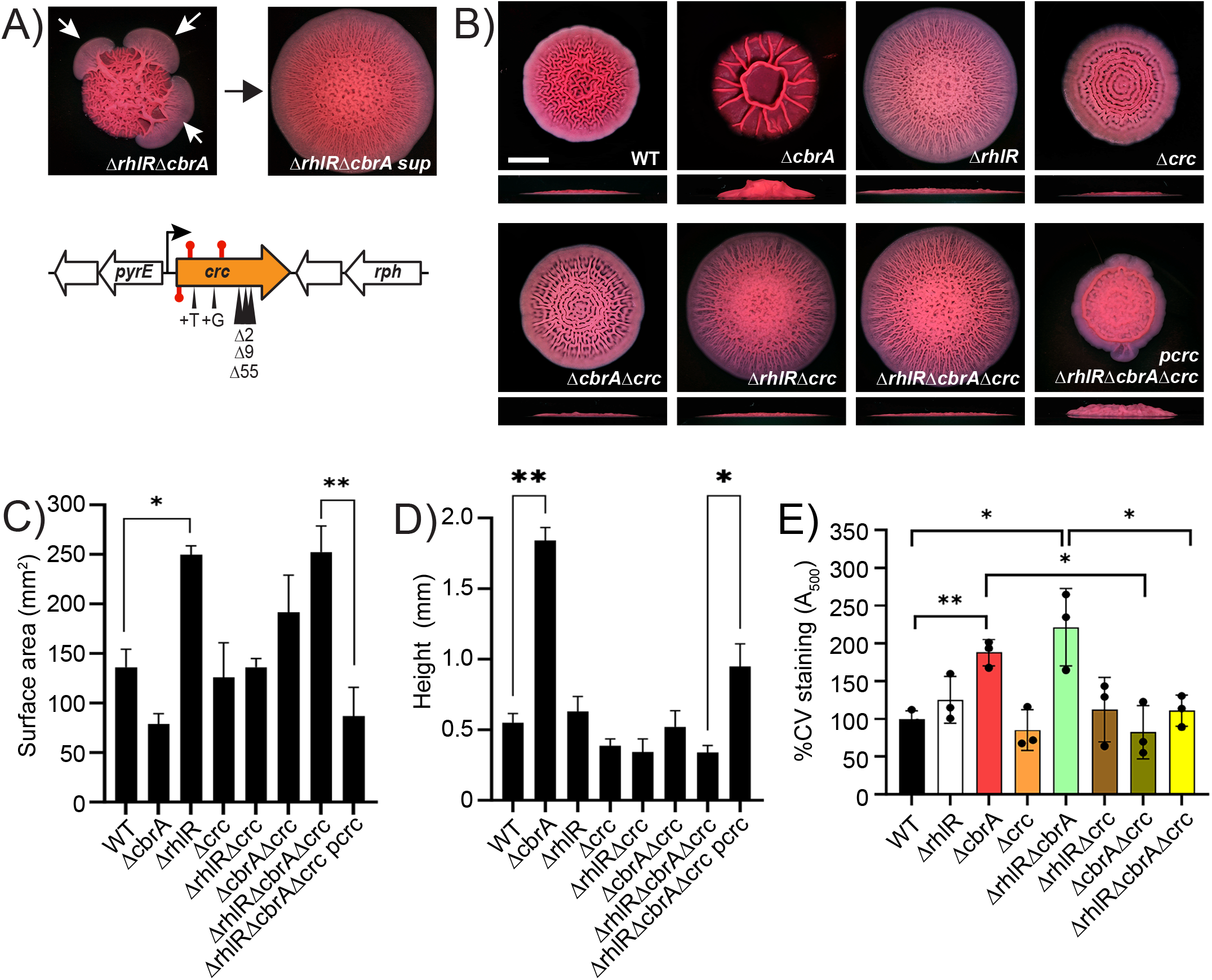
*P. aeruginosa* Δ*rhlR*Δ*cbrA* double mutant forms unstable biofilms. A) (Top) Shown is a representative isolation of a suppressor mutation of the Δ*rhlR*Δ*cbrA* biofilm phenotype. The white arrows in the left panel indicate flares radiating from the biofilm diagnostic of the emergence of suppressor mutations. The right panel shows the biofilm phenotype of a mutant following isolation. (Bottom) Chromosomal arrangement of the *crc* (orange) gene. Large white arrows represent open reading frames (lengths not to scale), black bent arrow indicates promoter, red stem-loops indicate STOP mutations and black triangles indicate the locations of insertion and deletion suppressor mutations. B) Colony biofilm phenotypes of WT PA14 and the designated mutants on Congo red agar medium after 120 h of growth. Scale bar, 5 mm. C) Colony biofilm surface area quantitation for the indicated strains after 120 h of growth. Error bars represent standard deviation of three independent experiments. Statistical significance was determined using unpaired *t*-test in GraphPad Prism software. ** P <0.01, * P <0.05. D) Colony biofilm height quantitation for the indicated strains after 120 h of growth. Error bars represent standard deviation of three independent experiments. Statistical significance was determined using two-tailed *t*-test with Welch’s correction in GraphPad Prism software. ** P <0.01, * P <0.05. E) Biofilm development assays followed by crystal violet staining for WT and indicated mutant strains. Error bars represent standard deviation of three biological replicates. Statistical significance was determined using two-tailed *t*-test with Welch’s correction in GraphPad Prism software. ** P <0.01, * P <0.05, and ns means not significant.

To explore the extent of biofilm regulation by Crc, we analyzed the ability of WT and the Δ*crc* single and combination mutants to form solid surface-associated (SSA) biofilms on borosilicate glass. Crystal violet staining of these SSA biofilms showed significant increase in mature biofilms in the Δ*cbrA* and Δ*rhlR*Δ*cbrA* mutants compared to WT (Fig. 2E). Absence of Crc significantly decreased SSA biofilms in both Δ*cbrA* and Δ*rhlR*Δ*cbrA* mutant backgrounds (Fig. 2E). Taken together, our data suggest a new role for Crc beyond reverse diauxie, i.e., Crc promotes biofilm development.

One known regulatory target of Crc beyond reverse diauxie is *phzM*, that encodes for a key enzyme involved in pyocyanin biosynthesis (Huang et al., 2012). Furthermore, the absence of phenazines contributes to the colony biofilm spreading phenotype of the Δ*rhlR* mutant (Mukherjee et al., 2017). To determine the contribution of phenazines to the Δ*rhlR*Δ*cbrA* double mutant phenotype, we deleted the two phenazine biosynthesis operons *phzA1-G1* and *phzA2-G2* (Δ*phz*) in the WT, the Δ*cbrA* single and the Δ*rhlR*Δ*cbrA* double mutant backgrounds and assayed pyocyanin production and biofilm development. As reported previously, the Δ*phz* mutant did not produce pyocyanin pigment and its biofilm spread radially outwards and covered more surface area than WT (Fig. S5A-C, Dietrich *et al*, 2013). Mutation of the phenazine biosynthesis operons, however, failed to increase biofilm spreading in cells lacking CbrA (Fig. S5B-C). Likewise, there was no detectable change in biofilm morphology of the Δ*rhlR*Δ*cbrA*Δ*phz* mutant when compared with its parent Δ*rhlR*Δ*cbrA* strain (Fig. S5B-C). We infer that phenazines do not affect biofilm morphology when CbrA is absent.

### CrcZ small RNA serves as the point of convergence for quorum and nutrient-sensing cues

To define the molecular basis underpinning the different Δ*rhlR* and Δ*cbrA* biofilm phenotypes, we used RNA-seq to compare the global transcriptional profiles of the biofilms of WT PA14 and the Δ*rhlR,* Δ*cbrA,* and Δ*rhlR*Δ*cbrA* double mutants. We harvested RNA from biofilms grown on Congo red agar medium for 72 h. Principal component analysis (PCA) of normalized read counts for 5,978 genes showed that biofilm samples of each mutant clustered separately from WT (Fig. S6). Comparative transcriptomic analysis revealed a total of 1366 differentially expressed genes (DEGs; genes with expression fold-changes ≤−2 and ≥2, and the P values (Padj), adjusted using the Benjamini-Hochberg procedure, <0.05; see Table S3 in the supplemental material) in the Δ*cbrA* mutant (Fig. 3A). In the Δ*rhlR* mutant, we find a total of 709 DEGs, of which 326 transcripts overlapped with the CbrA regulon. Comparing the Δ*rhlR*Δ*cbrA* double to the Δ*cbrA* single mutant shows a 51.6% reduction in the number of DEGs in the double mutant (Fig. 3A). Furthermore, 125 genes were uniquely regulated in the Δ*rhlR*Δ*cbrA* mutant (Fig. 3A). We conclude that the combined absence of each regulator – CbrA and RhlR – gives rise to a transcriptomic response distinct from mere addition of their individual regulons.

**Fig. 3:**
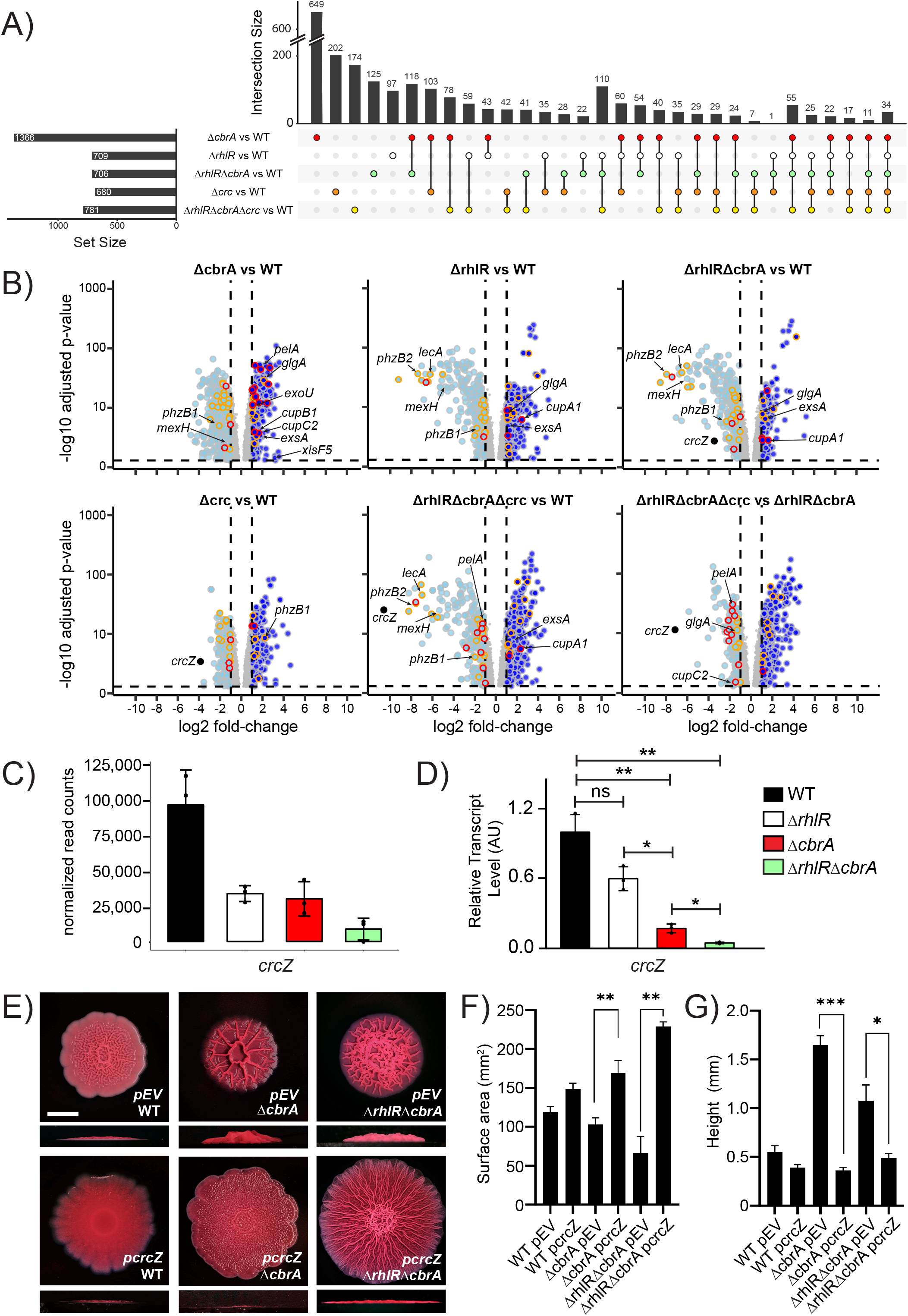
CrcZ small RNA allows integration of Rhl- and Cbr-signaling pathways and modulates biofilm development. A) Upset plot showing overlaps in genes that are differentially regulated in Δ*rhlR,* Δ*cbrA,* Δ*crc*, Δ*rhlR*Δ*cbrA* double, and Δ*rhlR*Δ*cbrA*Δ*crc* triple mutants compared to WT. Numbers on top of each vertical bar indicate number of genes differentially regulated in each intersection. Set size indicates the total number of genes that are significantly differentially regulated in a particular mutant when compared to WT. B) Volcano plots of RNA-seq data for Δ*rhlR,* Δ*cbrA,* Δ*crc*, Δ*rhlR*Δ*cbrA* and Δ*rhlR*Δ*cbrA*Δ*crc* compared to WT, and Δ*rhlR*Δ*cbrA*Δ*crc* compared to Δ*rhlR*Δ*cbrA*. Light blue solid circles with gray outlines represent genes with expression fold-changes ≤−2 and the P values (Padj), adjusted using the Benjamini-Hochberg procedure, <0.05; dark blue solid circles with gray outlines represent genes with expression fold-changes ≥2 and the P values (Padj), adjusted using the Benjamini-Hochberg procedure, <0.05. Gray solid circles represent genes with gene expression fold changes ≥-2 or ≤2 or Padj ≥ 0.05. Genes reported to be involved in virulence are outlined in yellow and those reported to be associated with biofilms are outlined in red. C) Normalized read counts obtained via Median Ratio Normalization (MRN) analysis for *crcZ* gene from RNAseq run on biofilm samples of WT and indicated mutants. D) Relative expression of *crcZ* gene normalized to 16S RNA, *ostA* and *rpsO* transcript levels measured by qRT-PCR in WT PA14 and indicated mutants after 120 h of colony biofilm growth. AU denotes arbitrary units. Error bars represent standard deviation of three biological replicates. Statistical significance was determined using two-tailed *t*-test with Welch’s correction in GraphPad Prism software. ** P <0.01, * P <0.05, and ns means not significant. E) Colony biofilm phenotypes of WT PA14 and the designated mutants on Congo red agar medium after 120 h of growth. Scale bar, 5 mm. F) Colony biofilm surface area quantitation for the indicated strains after 120 h of growth. Error bars represent standard deviation of three independent experiments. Statistical significance was determined using two-tailed *t*-test with Welch’s correction in GraphPad Prism software. ** P <0.01. G) Colony biofilm height quantitation for the indicated strains after 120 h of growth. Error bars represent standard deviation of three independent experiments. Statistical significance was determined using two-tailed *t*-test with Welch’s correction in GraphPad Prism software. *** P <0.001, ** P <0.01, * P <0.05.

Next, we determined the transcriptomic response to the absence of Crc. The Crc regulon consists of 680 genes, of which ∼48% are also regulated by CbrA and ∼36% coregulated by RhlR (Fig. 3A, Table S3). We note that, similar to the Δ*rhlR*Δ*cbrA* biofilm samples, a subset of genes was uniquely regulated in the Δ*rhlR*Δ*cbrA*Δ*crc* triple mutant, i.e., these 174 genes were not differentially expressed in the Δ*cbrA*, Δ*rhlR* and Δ*crc* single mutant strains compared to WT (Fig. 3A). Thus, the combined absence of RhlR, CbrA and Crc results in a distinct regulon that does not overlap with the single mutants. We infer that CbrA, RhlR and Crc regulons exhibit complex non-linear interactions.

The expression of the small RNA CrcZ that antagonizes Crc to allow coordinated utilization of carbon sources is activated by CbrA/CbrB (Fig. 1A; Sonnleitner et al., 2009; Sonnleitner et al., 2018). Thus, CrcZ levels are downregulated in the Δ*cbrA* mutant compared to WT (Fig. 3B). However, our RNA-seq data revealed a more severe reduction in CrcZ abundance in the Δ*rhlR*Δ*cbrA* double mutant compared to WT than in the Δ*rhlR* and Δ*cbrA* single mutants (Fig. 3B, 3C). Indeed, quantitative RT-PCR shows that *crcZ* transcript levels are ∼ 6-fold lower in the Δ*cbrA* mutant than WT but 23-fold lower in the Δ*rhlR*Δ*cbrA* mutant than WT after 120 h of colony biofilm growth (Fig. 3D). Furthermore, consistent with previous report that CrcZ small RNA is stabilized by Crc in *Pseudomonas putida* (Hernandez-Arranz et al., 2016), we find significant reduction in CrcZ abundance in the Δ*crc* and Δ*rhlR*Δ*cbrA*Δ*crc* mutants compared to WT PA14 (Fig. 3A-B, Table S3). Taken together, our data suggest that low levels of CrcZ promote biofilm morphology with reduced surface coverage and increased height.

To assess the contribution of CrcZ small RNA to biofilm development, we introduced a plasmid expressing *crcZ* from *Plac* promoter to WT and the Δ*cbrA* single and Δ*rhlR*Δ*cbrA* double mutants. Overexpression of CrcZ small RNA led to increased surface area coverage and a severe reduction in biofilm height in the Δ*cbrA* mutant compared to the empty vector control (Fig. 3E-G). Furthermore, overexpression of CrcZ small RNA in the Δ*rhlR*Δ*cbrA* double mutant resulted in a hyper-rugose biofilm that resembled the Δ*rhlR* mutant in terms of biofilm morphology, surface area and height (Fig. 3E-G). We conclude that the abundance of CrcZ small RNA is a key determinant of biofilm development in response to signals transduced via CbrA and RhlR. We further conclude, that reduced *crcZ* expression leads to high Crc activity which in turn restricts surface area coverage and promotes height, likely explaining why *crc* suppressors arose in the Δ*rhlR*Δ*cbrA* mutant to allow biofilm expansion.

### Crc activates the expression of biofilm matrix components

Closer inspection of the transcriptomics data revealed the over expression of genes involved in Pel biosynthesis such as *pelA* in the Δ*rhlR*Δ*cbrA* double mutant compared to the Δ*rhlR*Δ*cbrA*Δ*crc* triple mutant (Fig. 3B and Table S3). These data suggest that Crc can be a positive regulator of Pel biosynthesis. To validate the role of Crc in activating *pelABCDEFG* operon expression, we performed quantitative RT-PCR for *pelA* from colony biofilm samples in the WT and a series of mutants - Δ*rhlR,* Δ*cbrA,* Δ*crc,* Δ*rhlR*Δ*cbrA,* Δ*rhlR*Δ*crc,* Δ*cbrA*Δ*crc* and Δ*rhlR*Δ*cbrA*Δ*crc* (Fig. 4A, S7A-B). We performed these experiments on biofilms grown for 72 h i.e., early biofilms and 120 h i.e., mature biofilms. Our analyses demonstrate that RhlR activates the expression of *pelA* as the Δ*rhlR* mutant showed four-fold reduction in *pelA* transcript levels compared to WT in mature biofilms (Fig. 4A). Thus, although RhlR represses biofilm surface area coverage, it is a positive regulator of Pel biosynthesis.

**Fig. 4:**
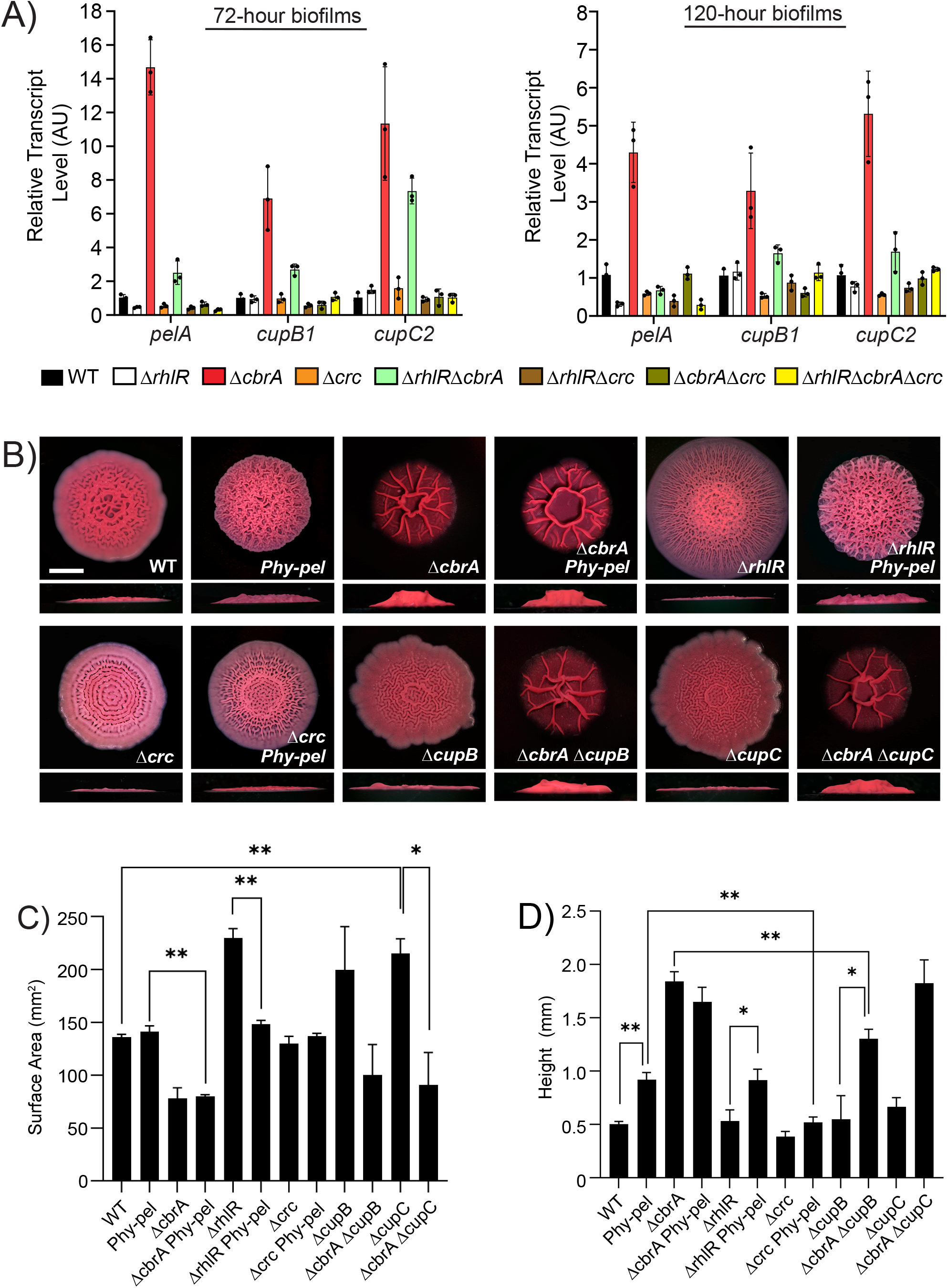
Crc promotes biofilm matrix gene expression. A) Relative expression of *pelA*, *cupB1*, *cupC2* genes normalized to 16S RNA, *ostA* and *rpsO* transcript levels in WT PA14 and indicated mutants after 72 h and 120 h of colony biofilm growth. AU denotes arbitrary units. Error bars represent standard deviation of three biological replicates. B) Colony biofilm phenotypes of WT PA14 and the designated mutants on Congo red agar medium after 120 h of growth. Scale bar, 5 mm. C) Colony biofilm surface area quantitation for the indicated strains after 120 h of growth. Error bars represent standard deviation of three independent experiments. Statistical significance was determined using two-tailed *t*-test with Welch’s correction in GraphPad Prism software. ** P <0.01, * P <0.05. D) Colony biofilm height quantitation for the indicated strains after 120 h of growth. Error bars represent standard deviation of three independent experiments. Statistical significance was determined using two-tailed *t*-test with Welch’s correction in GraphPad Prism software. ** P <0.01, * P <0.05.

Conversely, we find a fifteen-fold and five-fold increase in *pelA* transcript levels in the Δ*cbrA* mutant compared to WT at 72 h and 120 h timepoints, respectively. In addition, there is a two-fold reduction in the relative abundance of *pelA* transcript in the Δ*crc* mutant compared to WT at 72 h and 120 h timepoints respectively, and twentyseven-fold and eight-fold reduction compared to the Δ*cbrA* strain at 72 h and 120 h timepoints respectively. Furthermore, the absence of Crc is epistatic to Δ*cbrA* as *pelA* transcript levels are drastically reduced in the Δ*cbrA*Δ*crc* and Δ*rhlR*Δ*cbrA*Δ*crc* mutants (Fig. 4A). Thus, we conclude that Crc activates *pelABCDEFG* operon expression and the high expression of *pelABCDEFG* operon in the Δ*cbrA* and Δ*rhlR*Δ*cbrA* mutants is due to the unchecked activity of Crc in the absence of CrcZ small RNA.

To determine whether different expression level of *pelABCDEFG* operon is involved in the distinct surface area and height of the Δ*rhlR* and Δ*cbrA* mutants, we generated a Pel overexpression strain where we replaced the native promoter of *pelABCDEFG* operon with a constitutively expressed artificial promoter (*PpelA::Physpank*; we call this strain *Phy-pel*). Compared to WT, relative transcript abundance of *pelA* is increased about eighteen folds in the the *Phy-pel* strain (Fig. S7B). Accordingly, the *Phy-pel* strain exhibits a hyper-wrinkled biofilm phenotype that is distinct from WT in the colony biofilm assay (Fig. 4B), and increased attachment to borosilicate glass in the solid-surface attachment assay (Fig. 5A). Consistent with the result that RhlR activates the expression of *pelA* (Fig. 4A), overexpression of *pelABCDEFG* is epistatic to Δ*rhlR* as the Δ*rhlR Phy-pel* strain shows reduced biofilm spreading compared to the parent Δ*rhlR* mutant (Fig. 4B-C). Furthermore, the absence of *cbrA* is epistatic to overexpression of *pelABCDEFG* as the Δ*cbrA Phy-pel* double mutant formed biofilms identical to the Δ*cbrA* mutant (Fig. 4B-D). Although the abundance of *pelA* transcript is similar between Δ*cbrA* and *Phy-pel* mutants, the *Phy-pel* strain produced a hyper-rugose biofilm morphology that is markedly distinct from the Δ*cbrA* mutant biofilm (Fig. 4B-D, S7B). We infer (and we return to this point later) that the hyper-rugose biofilms of Δ*cbrA* mutant likely involves another biofilm matrix factor in addition to Pel exopolysaccharide. Notably, the Δ*crc Phy-pel* double mutant mirrors the Δ*crc* mutant biofilm phenotypes in the colony biofilm and solid-surface attachment assays (Fig. 4B-D, 5A). Thus, we conclude that the absence of *crc* is epistatic to transcriptional activation of the *pelABCDEFG* operon.

**Fig. 5:**
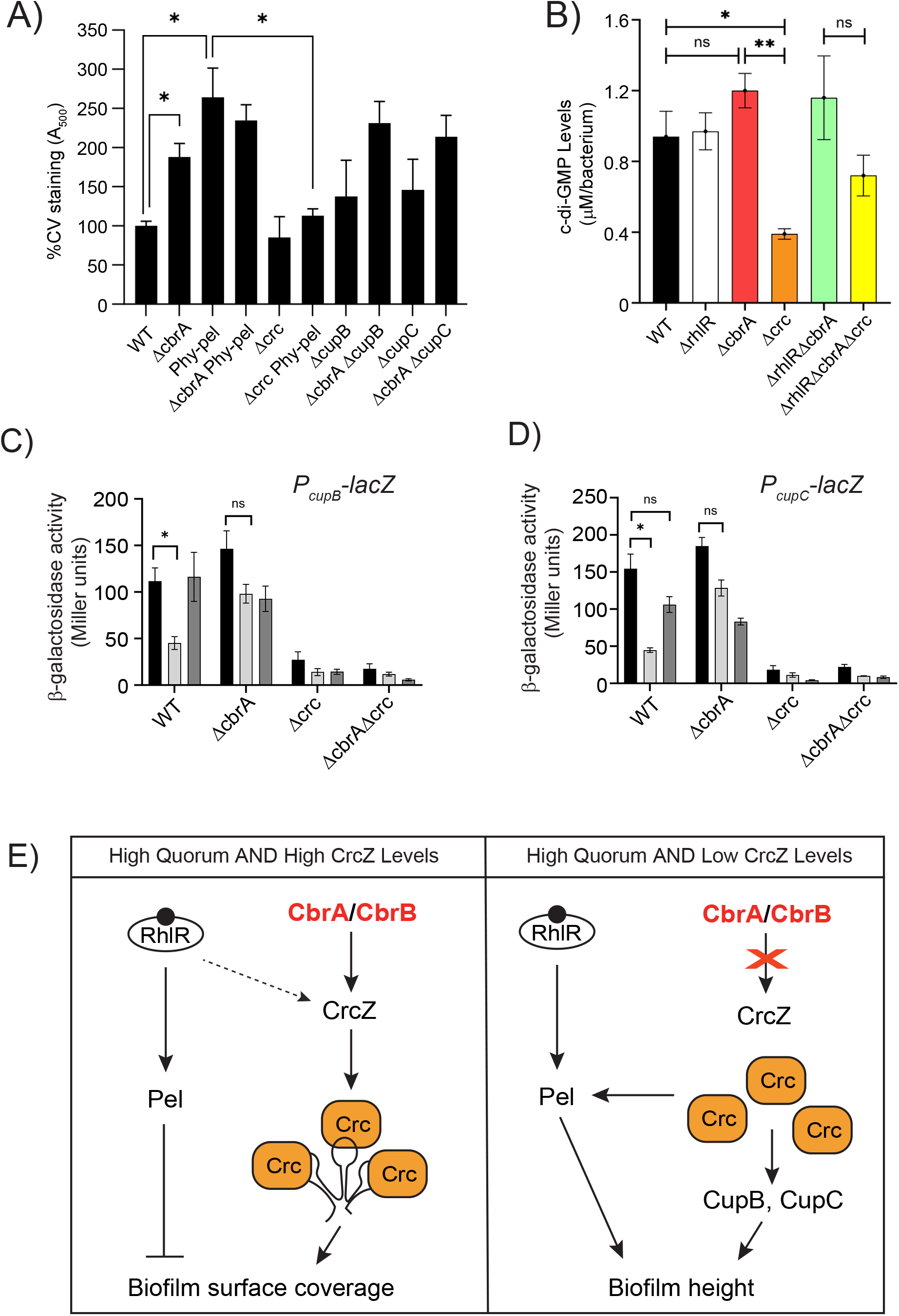
Crc dependent increase in *pel* expression is post-transcriptional. A) Biofilm crystal violet staining assays for WT and indicated mutant strains. Error bars represent standard deviation of three biological replicates. Statistical significance was determined using two-tailed *t*-test with Welch’s correction in GraphPad Prism software. * P <0.05. B) c-di-GMP levels of the indicated strains. Error bars represent standard deviation of three biological replicates. Statistical significance was determined using unpaired *t*-test in GraphPad Prism software. ** P <0.01, * P <0.05, and ns means not significant. C-D) β-galactosidase assays of *PcupB-lacZ* transcriptional (B) or *PcupC-lacZ* transcriptional (C) fusions for background genotypes indicated on the X-axis and grown in tryptone broth with no addition (black), 50mM pyruvate (light grey), or 50mM glucose (dark grey). Error bars represent SEM of three biological replicates. Statistical significance was determined using two-tailed *t*-test with Welch’s correction in GraphPad Prism software. * P <0.05, and ns means not significant. C) Model for Crc-dependent activation of biofilm matrix components. At high population density, RhlR mediated quorum sensing activates the expression of Pel exopolysaccharide while Crc, depending on the abundance of CrcZ small RNA, promotes the expression of Pel, CupB and CupC fimbriae components of the biofilm matrix. Arrow indicates activation and T-bar indicates inhibition.

Because Crc functions as a post-transcriptional regulator in reverse carbon catabolite repression, we hypothesized that Crc mediates increase in *pelABCDEFG* expression via post-transcriptional mechanism. One well-known post-transcriptional mode of control of Pel biosynthesis is the small molecule c-di-GMP such that high c-di-GMP levels correlate with biofilm formation (reviewed in Ha and O’Toole, 2015). Furthermore, Hfq, and thereby carbon catabolite repression, has been linked to c-di-GMP concentration in cells (Pusic, et al., 2018). This prompted us to test whether changes in intracellular c-di-GMP concentration played a role in the difference in biofilm development between the mutants in this study. Fig. 5B shows that while c-di-GMP concentration is lowered in the Δ*crc* mutant compared to WT, none of the Δ*rhlR,* Δ*cbrA,* Δ*rhlR*Δ*cbrA* and Δ*rhlR*Δ*cbrA*Δ*crc* mutants displayed any discernable changes from WT. We conclude that c-di-GMP levels do not contribute to the difference in biofilm development of the Δ*rhlR,* Δ*cbrA,* and Δ*rhlR*Δ*cbrA* mutants.

In addition to Pel, the biofilm matrix involves a large arsenal of cell surface-associated structures, including flagella, type IVa or type IVb pili, Fap fibrils and the cup fimbriae (Ruer et al. 2007; de Bentzmann et al. 2006; Klausen et al. 2003; O’Toole and Kolter, 1998; Vallet et al. 2001; Dueholm et al. 2010). *P. aeruginosa* can produce five types of cup fimbriae – CupA, CupB, CupC, CupD and CupE, products of *cupA*, *cupB*, *cupC*, *cupD*, and *cupE* gene clusters each of which encodes an usher, a chaperone, and at least one fimbrial subunit. Our RNA-seq data revealed that genes required for the biosynthesis of CupB and CupC matrix components are upregulated in the Δ*cbrA* and Δ*rhlR*Δ*cbrA* mutants while those for CupA are upregulated in the Δ*rhlR* and Δ*rhlR*Δ*cbrA* mutants (Fig. 3B, Table S3). To assess the coordinated regulation of the Cup components of the biofilm matrix by RhlR, CbrA and Crc over time, we performed quantitative RT-PCR for *cupA1*, *cupB1*, *cupC2*, *cupD4*, and *cupE1* from colony biofilm samples after 72 and 120 hours of growth, respectively, in the WT and the Δ*rhlR,* Δ*cbrA,* Δ*crc,* Δ*rhlR*Δ*cbrA,* Δ*rhlR*Δ*crc,* Δ*cbrA*Δ*crc* and Δ*rhlR*Δ*cbrA*Δ*crc* mutants (Fig. 4A, S7A). Our data show that the relative mRNA levels of the *cupB1* and *cupC2* genes increased greater than five-folds at 72 h and more than three-folds at 120 h in the Δ*cbrA* single and Δ*rhlR*Δ*cbrA* double mutants but these increases in transcript levels were abolished in the Δ*cbrA*Δ*crc* double and Δ*rhlR*Δ*cbrA*Δ*crc* triple mutants (Fig. 4A, S7A). This suggests that, in addition to Pel, Crc promotes the expression of CupB and CupC matrix components.

The CupB and CupC fimbriae could contribute to increased adhesive properties of the Δ*cbrA* biofilm due to the high expression of *cupB* and *cupC* operons in this mutant (Fig. 4A; Vallet et al, 2001). To determine the role of CupB and CupC fimbriae in biofilms of the Δ*cbrA* mutant, we generated deletion mutants where we removed entire *cupB* (*cupB1-B5*) and *cupC* (*cupC1-C3*) operons in WT and Δ*cbrA* backgrounds. The Δ*cbrA*Δ*cupB* mutant exhibited significant, albeit minor, decrease in biofilm height compared to the Δ*cbrA* mutant while for the Δ*cbrA*Δ*cupC* mutant, surface area coverage and height parameters show values similar to the Δ*cbrA* mutant (Fig. 4B-D). We infer that although Crc activates the expression of both *cupB* and *cupC* operons, only CupB appears to contribute to biofilm matrix in laboratory settings.

Next, to probe the expression of biofilm matrix genes in response to carbon sources we made transcriptional reporter fusions to the *cupB* and *cupC* promoters (P*cupB-lacZ* and P*cupC-lacZ*). We incorporated the reporter fusion into an intergenic region on the chromosomes of WT *P. aeruginosa* and the Δ*cbrA,* Δ*crc* and Δ*cbrA*Δ*crc* mutants. The P*cupB-lacZ* and P*cupC-lacZ* reporters exhibited approximately four-fold and eight-fold lower expression in the Δ*crc* and Δ*cbrA*Δ*crc* mutants than the WT in tryptone broth (Fig. 5C-D). These results show that Crc is absolutely required for the expression of *cupB* and *cupC* operons in both WT and Δ*cbrA* backgrounds. In *P. aeruginosa*, “reverse” carbon catabolite repression is known to be triggered by organic acids such as pyruvate (Collier et al., 1996; Liu, 1952; MacGregor et al., 1991; Wolff et al., 1991). Furthermore, pyruvate activates the expression of *crcZ* in both *P. aeruginosa* and *P. putida* (Valentini et al, 2014). Therefore, we tested pyruvate as preferred carbon source and glucose as non-preferred sugar. Expression of the reporters in the WT was reproducibly and significantly lowered in the presence of pyruvate but not glucose (Fig. 5C-D). Taken together, our data lead to a regulatory model in which environmental cues that reduce the levels of CrcZ small RNA allow for Crc-dependent activation of biofilm matrix gene expression in *P. aeruginosa* (Fig. 5E).

Bacteria of the genus Pseudomonas are metabolically versatile, can be isolated from a wide range of niches and preferentially catabolize organic acids as they lack a functional glycolytic pathway (reviewed in Rojo, 2010). This “reverse” carbon catabolite repression is thought to provide adaptive advantage in different niche colonization. As such, the CbrA/CbrB/CrcZ/Crc signaling cascade is conserved in most Pseudomonads and components of this pathway have been found to be exchangeable between *P. aeruginosa* and *P. putida* (Valentini et al., 2014). We explored whether Crc in other species of *Pseudomonas* might also be involved in controlling biofilm formation via phylogenetic analysis of the co-occurrence of *cbrA, crc, pelA, cupB1* and *cupC2* in the genomes of diverse Pseudomonads (Fig. 6). Consistent with previous reports, majority of the Pseudomonads encode for CbrA and Crc orthologs. PelA and CupB1 are found to co-occur in a subset of the genomes that include several CF isolates of *P. aeruginosa* while CupC2 is present in ∼45% of the genomes analyzed here. For example, Pseudomonads that colonize the rhizosphere of plants such as *P. stutzeri*, *P. chlororaphis*, *P. syringae* and *P. protegens* encode for genes required to assemble CupC fimbriae. These bacteria are exposed to glucose as well as organic acids like succinate and pyruvate that are commonly found in root exudates and likely derive ecological advantage by integrating nutritional cues to control adhesion to root surfaces. Thus, it appears that Crc has a conserved function in the regulation of CupC fimbriae which might also contribute to biofilm development in the environmental Pseudomonads.

**Fig. 6:**
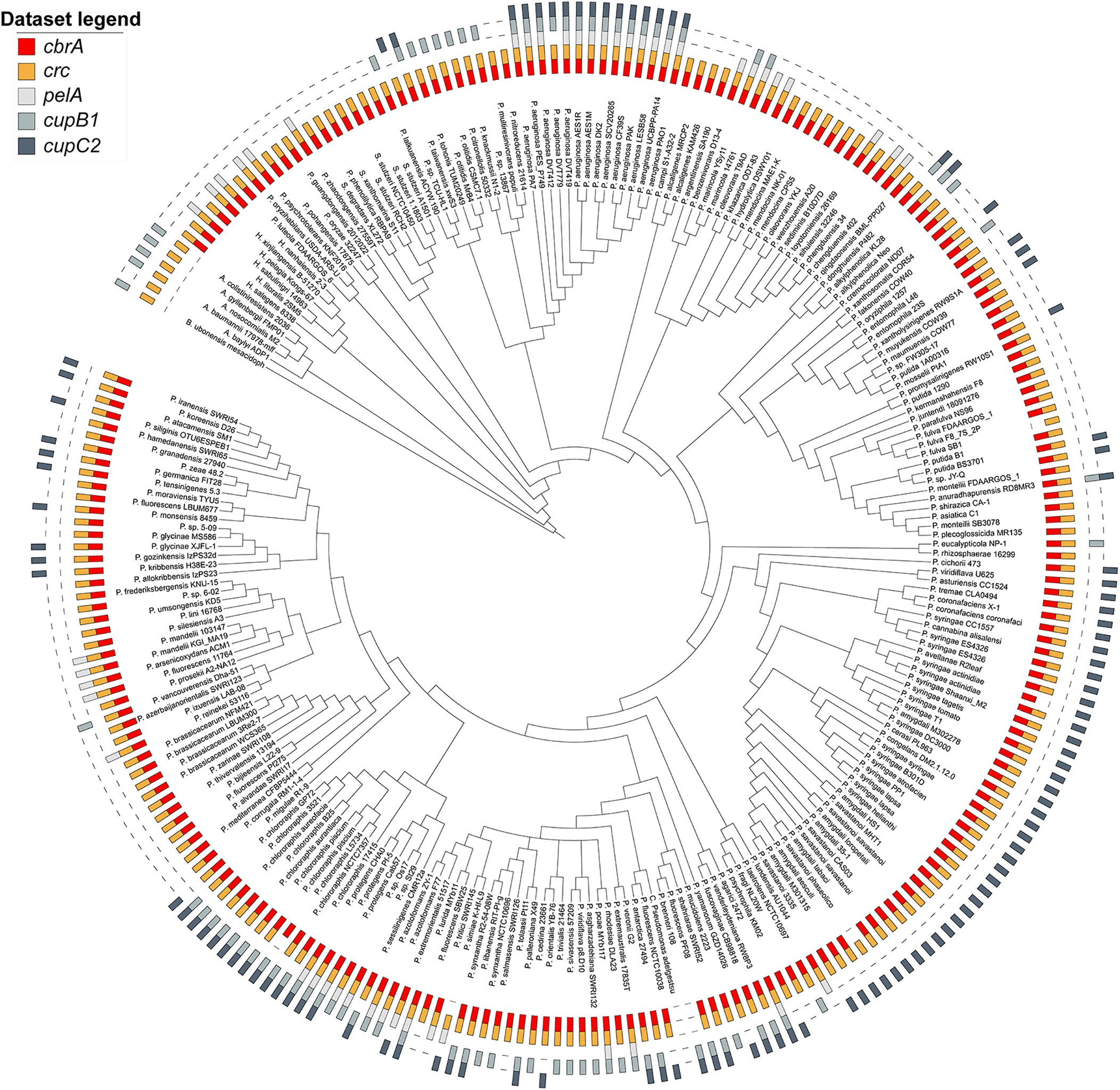
The CbrA and Crc proteins and their regulatory targets Pel, CupB and CupC are conserved in diverse Pseudomonads. Co-occurrences of CbrA and Crc are depicted in red and orange, respectively, while that of their regulatory target genes obtained from this study – *pelA*, *cupB1*, and *cupC2* - are depicted in shades of grey as indicated in the dataset legend.

## DISCUSSION

Combinatorial gene regulation by multiple two-component systems is highly utilized in bacteria in responding to environmental cues to allow distinct gene expression patterns. *P. aeruginosa* is a versatile human pathogen that causes both acute and chronic infections such as in the lungs of cystic fibrosis (CF) patients and in patients suffering from burn and diabetic wounds that are often associated with worse disease outcomes (Rumbaugh et al., 1999; Hoffman et al, 2009; Hammond et al, 2015; Feltner et al, 2016). In this study, we sought to define the interaction of Rhl quorum sensing with another global regulatory system, i.e., CbrA/CbrB pathway. As the sensory signal that activates CbrA kinase is unknown (reviewed in Monteagudo-Cascales et al. 2022), we studied biofilm development and biofilm-associated transcriptomic changes in the absence of RhlR, CbrA and both. Our data highlight the complexity of sensory regulation mediated by these two signaling pathways in *P. aeruginosa* biofilms and reveals a previously unexplored facet of Crc as it promotes expression of specific biofilm matrix components – Pel exopolysaccharide and CupB/CupC fimbriae.

*P. aeruginosa*, one of the major causes of morbidity and mortality of CF patients, colonize the lung mucus which is a complex substrate that provides carbon, nitrogen and other energy sources during infection (Palmer et al, 2005). Although glucose is a less preferred carbon source for *P. aeruginosa*, it is present at high concentration in the mucus of both CF and non-CF patients, with CF mucus typically having higher glucose content than those of other pulmonary diseases (Palmer et al, 2007; La Rosa et al, 2019). Our work suggests that glucose (or another less preferred carbon source) could function as an inducer of biofilm formation via the CbrA/CbrB/CrcZ/Crc pathway. We do not yet know the molecular mechanism by which Crc activates the expression of *pel, cupB* or *cupC* genes. We speculate that one possible mechanism might be post-transcriptional for *pel* as the absence of Crc decreases biofilm formation even after exchanging the *PpelA* promoter with an artificial promoter. Nonetheless, given that both carbon catabolite repression and Pel biosynthetic genes are widespread in bacteria (Whitfield, 2021), we expect future studies defining the molecular details underlying the link between “reverse” carbon catabolite repression and *pel* expression would be informative.

The material properties of the matrix contribute to biofilm growth and architecture. For instance, in *Vibrio cholerae* and *Bacillus subtilis*, matrix production allows biofilm-dwelling cells to establish an osmotic pressure difference between the biofilm interior and the external environment that promotes colony biofilm expansion (Yan et al, 2017; Seminara et al., 2012). Specifically, *V. cholerae* colony biofilms were found to expand more and cover greater surface area in the absence of the matrix protein RbmA (Yan et al., 2017). Our work suggests that in PA14, lower Pel polysaccharide levels allow greater surface area coverage as observed in the Δ*rhlR* mutant. Conversely, increased Pel polysaccharide levels limits biofilm expansion as is the case with Δ*cbrA* and Δ*rhlR* Δ*cbrA* mutants.

*P. aeruginosa* PA14 has five different *cup* operons that encode proteins involved in the assembly of surface adhesive fimbriae, but the *cup* genes are poorly expressed in standard laboratory conditions making it difficult to study their regulation. To date, few regulators such as the H-NS-like protein MvaT that represses *cupA* gene expression and the two-component systems named Roc1 and Roc2 that induce the expression of *cupB* and *cupC* genes have been described (Vallet-Gely et al., 2005; Ruer et al., 2007). Nonetheless, the Cup fimbrial structures have been reported to facilitate bacterial attachment to host tissue, and promote biofilm formation and pathogenesis (Busch et al., 2012; Vallet et al., 2001). Our finding that organic acids such as pyruvate can repress the expression of *cupB* and *cupC* fimbrial genes hints at change in adhesion depending on nutritional cues that might contribute to biofilm architecture within hosts.

Transcriptome analysis may reveal both direct and indirect targets of Crc. Crc/Hfq binding to transcripts is thought to involve at least one CA motif (AAnAAnAA) in the neighborhood of the translational start site in target mRNAs (Sonnleitner E, et al. 2018, Kambara et al., 2018). Accordingly, we scanned the 5’ untranslated region upstream of the genes that are regulated by Crc in two RNA-seq datasets - Δ*crc* versus WT and Δ*rhlR*Δ*cbrA* versus Δ*rhlR*Δ*cbrA*Δ*crc* mutants. We found 32 and 73 Crc-regulated transcripts having a putative CA motif respectively, but none of these transcripts belong to biofilm-matrix biosynthesis genes (Table S4). One caveat is that the Crc/Hfq recognition motif for biofilm-matrix biosynthesis genes might be distinct from the aforementioned CA motif. Another possibility is that Crc indirectly activates the expression of biofilm matrix components.

Our RNA-seq analysis sheds light on the transcriptomic landscape of WT *P. aeruginosa* biofilms and provides a map of the regulatory network orchestrated by RhlR, CbrA and Crc. When we bin the differentially expressed genes according to their function defined by their PseudoCAP categories (Winsor et al., 2016), CbrA appears to upregulate the “Carbon compound metabolism”, “Amino acid biosynthesis and metabolism”, “Central intermediary metabolism” and “Energy metabolism” functional classes (Fig. S8). In addition, CbrA controls the expression of several genes in the “Biosynthesis of cofactors, prosthetic groups, “Two component regulatory system”, “Transport of small molecules”, “Protein secretion/export” and “Membrane proteins” functional classes. For instance, genes involved in the Type II secretion system (*xcpUVWYZ* operon) and the Hcp Secretion Island-I-encoded type VI secretion system (H1-T6SS) are activated while those involved in Type III secretion system (*exsB, exsC, pcrV*) are repressed by the Cbr system (Table S3). This highlights the importance of the Cbr system in cellular physiology in biofilms beyond carbohydrate and amino acid metabolism.

The RhlR regulon exhibits enrichment in the “Secreted factors”, “Chaperones and heat shock proteins” and “Adaptation/Protection” functional classes (Fig. 3B). However, most of genes regulated by CbrA and RhlR belong to the “Hypothetical, unclassified, unknown” category limiting the power of our analysis. Nonetheless, we note that several virulence factors including *lecA*, *phz1*, *phz2*, *mexH* are downregulated in the Δ*rhlR* and Δ*rhlR*Δ*cbrA* mutants, highlighting the importance of RhlR in *P. aeruginosa* virulence (Fig. 3B). Comparing the Δ*rhlR*Δ*cbrA* and Δ*rhlR*Δ*cbrA*Δ*crc* mutants shows that Crc represses several genes in the “Transport of small molecules”, “Energy metabolism”, and “Relative to phage, transposon or plasmid” classes (Fig. S8). Notably, in biofilms, Crc appears to both positively and negatively regulate several categories unrelated to carbon catabolite repression such as “Cell wall/LPS/Capsule”, “Protein secretion/export apparatus” and “Adaptation, Protection” (Fig. S8). Therefore, by characterizing the convergence of global signaling pathways on biofilm formation, we have expanded our understanding of the molecular interplay between key regulators - CbrA and RhlR, and identified Crc as a master regulator of biofilm matrix gene expression in response to environmental stimuli..

## MATERIALS AND METHODS

### Strains and growth conditions

*P. aeruginosa* UCBPP-PA14 strain was grown in lysogeny broth (LB) (10 g tryptone, 5 g yeast extract, 5 g NaCl per L), in 1% Tryptone broth (TB) (10 g tryptone per L) and on LB plates fortified with 1.5% Bacto agar at 37°C. When appropriate, antimicrobials were included at the following concentrations: 400 µg/mL carbenicillin, 50 µg/mL gentamycin, 100 μg/mL irgasan.

### Strain construction

Strains and plasmids were constructed as described previously (Mukherjee et al. 2017). To construct marker-less in-frame chromosomal deletions in *P. aeruginosa*, DNA fragments flanking the gene of interest were amplified, assembled by the Gibson method, and cloned into pEXG2 (Hmelo et al., 2015). The resulting plasmids were used to transform *Escherichia coli* SM10λ*pir*, and subsequently, mobilized into *P. aeruginosa* PA14 via biparental mating. Exconjugants were selected on LB containing gentamicin and irgasan, followed by recovery of deletion mutants on LB medium containing 5% sucrose. Candidate mutants were confirmed by PCR. The *cbrA* complementation plasmid was constructed by inserting DNA containing the promoter of *dksA* and the entire *cbrA* open-reading frame using HindIII and XbaI, followed by cloning into similarly digested pUCP18. The *crc* complementation plasmid was constructed by inserting DNA containing the promoter and entire *crc* open-reading frame using HindIII and XbaI, followed by cloning into similarly digested pUCP18. The *crcZ* overexpression plasmid was constructed by inserting DNA containing the entire *crcZ* gene using HindIII and XbaI, followed by cloning into similarly digested pUCP18.

To construct the P*cupB*-*lacZ* and P*cupC*-*lacZ* transcriptional reporter fusions, 500 bp of DNA upstream of the *cupB1 and cupC1* genes and the DNA encoding the *lacZ* open-reading frame were amplified using *P. aeruginosa* PA14 genomic DNA and the plasmid pIT2 as templates, respectively. Next, two DNA fragments of ∼730 bp, one corresponding to the intergenic region ∼700 bp downstream of the *P. aeruginosa PA14_20500* gene and the other corresponding to ∼1000 bp upstream of the *P. aeruginosa PA14_20510* gene, were amplified from *P. aeruginosa* PA14 genomic DNA. The four DNA fragments were assembled by the Gibson method and cloned into pEXG2. The resulting plasmid was used to transform *E. coli* SM10λ*pir*, and subsequently mobilized into *P. aeruginosa* PA14 WT and the Δ*rhlR* and Δ*rhlI* mutants via biparental mating as described above.

### Colony biofilm assay

One microliter of overnight *P. aeruginosa* cultures grown at 37°C in 1% Tryptone broth was spotted onto 60 x 15 mm Petri plates containing 10 mL 1% Tryptone medium fortified with 40 mg per L Congo red and 20 mg per L Coomassie brilliant blue dyes, and solidified with 1% agar. Colonies were grown at 25°C and images were acquired after 120 h using a Zeiss AxioZoom v16 microscope.

### Pellicle biofilm assay and crystal violet staining of SSA biofilms

*P. aeruginosa* was cultured overnight in 200 µL LB in a BioTek Synergy Neo2 microplate reader at 37°C under shaking conditions. The OD600 of the culture was taken by the plate reader to calculate the amount of culture needed to inoculate 2 ml 1% Tryptone broth so that the final OD_600_ is 0.005. Pellicle biofilms were developed for 72 hours in stationary 16*100 mm glass test tubes at room temperature. After 72 h, the cultures were vortexed to disrupt the pellicles. The samples were washed vigorously with tap water and were left to dry for an hour. The biofilms were then stained with 4 mL 0.1% crystal violet solution for half hour. The excess solution was poured out, the samples were washed thrice with tap water and let to dry overnight. For elution, 4 mL of 33% glacial acetic acid solution was used in each tube and samples were left to rest for an hour. The crystal violet stain was quantified at OD_550_ using a spectrophotometer (Thermo Scientific Genesys 20).

### RNA-seq

*P. aeruginosa* colony biofilms were grown for 40 h before harvested with a plastic inoculation loop. Half of each biofilm was picked up and transferred into 600 µL of Tri-reagent (Zymo Research) and disrupted by pipetting with a 1 mL tip. Total RNA was extracted with the Zymo Direct-zol RNA kit following the manufacturer’s instructions. Samples were subjected to DNAse treatment using Ambion DNase I (RNase-free) kit, followed by rRNA depletion and library preparation with Illumina Stranded Total RNA Prep with Ribo-Zero kit. Libraries were sequenced on Illumina’s NextSeq 2000, with 51 bp paired-end reads and 10 bp long index reads (51-10-10-51).

### RNA-seq analysis

RNA-seq reads were processed using the nf-core RNAseq pipeline version 3.4. Briefly, low-quality bases and contaminant adapters were trimmed using Cutadapt version 3.4 and Trimgalore version 0.6.7. RNA-seq reads passing quality filters were mapped against the reference genome of *P. aeruginosa* UCBPP_PA14 strain (GenBank assembly accession number CP000438.1) using HISAT2 version 2.2.0 (with parameters --rna-strandness RF, --no-mixed, --no-discordant, --no-spliced-alignment; Kim et al., 2019). Next, read mappings for each annotated gene were counted using the featureCounts program within the Subread package version 2.0.1 (with parameters -B -C and -s 2, Liao et al. 2014). Analysis of differentially expressed genes (DEGs) was performed using the DESeq2 package 1.28.0 (Love et al. 2014). Genes were considered significantly differentially expressed when the P value (Padj), adjusted using the Benjamini-Hochberg procedure, was <0.05. Assessment of functional classification enrichment was performed by assigning DEGs to at least one of 27 manually-defined and curated PseuodCAP functional classifications (Winsor et al., 2005). The percentage of genes in each category that exhibited downregulation or upregulation was then calculated.

### qRT-PCR

*P. aeruginosa* colony biofilms were grown for 40 h before harvested with a plastic inoculation loop. Half of each biofilm was picked up and transferred into 600 ul of Tri-reagent (Zymo Research) and disrupted by pipetting with a 1 ml tip. Total RNA was extracted with the Zymo Direct-zol RNA kit following the manufacturer’s instructions and quantified with a BioTek Synergy Neo2 microplate reader. 0.5 μg of total RNA was used for reverse-transcription using the TaKaRa PrimeScript RT Reagent Kit with gDNA Eraser. The resulting cDNA was used for real-time PCR using the Applied biosystems PowerTrack SYBR Green Master Mix on a Bio-Rad C1000 Touch Thermal Cycler. The results were exported into RDML v1.1 format and analyzed with web-based LinRegPCR (https://www.gear-genomics.com/rdml-tools/). Relative fold change was calculated with the qBase method (Hellemans et al. 2007) using *16s, clpX*, *ostA* and *rpsO* as reference genes.

### β-galactosidase assay

Briefly, bacterial cultures were grown to OD600 1.3. Pellets of 1mL cultures were collected and redissolved in 1mL Z-buffer with the addition of 200μg lysozyme for permeabilization. Samples were then incubated in 30°C for 15 min and diluted as needed in a total volume of 500μL. 4mg ONPG solution was added in the samples and time was recorded. The samples were incubating in 30°C until sufficient color change was observed. The reaction was quenched by 250μL 1M Na_2_CO_3_. Standard activity was calculated in Miller units.

### Intracellular cyclic di-GMP measurements

For intracellular cyclic di-GMP determination, cells were grown in tryptone broth and an aliquot was taken for OD measurement. 1 mL of cells were spun down, resuspended in 100 µL nucleotide extraction solution (40% acetonitrile, 40% Methanol, 0.1% Formic acid and 19.9% Water) and incubated at −20°C for 20 min. The samples were centrifuged at 15000 rpm for 5 min and the supernatant was transferred to a fresh tube. The samples were dried under vacuum, resuspended in 100 µL HPLC grade H2O and analyzed by HPLC-MS/MS as described previously (Massie et al, 2012). Intracellular cyclic di-GMP levels was determined by fitting the peak intensity to a standard curve with the known concentrations of cyclic di-GMP and by normalizing the levels of to the total number of cells.

### Phylogenetic tree construction

Genomes and associated proteomes GFF annotation files of strains from the Pseudomonas (238), Acinetobacter (5), and Burkholderia (1) genera were downloaded from the NCBI datasets database. To determine orthologous relationships between protein-coding genes, we used OrthoFinder version 2.5.4. The analysis was performed on an AWS EC2 instance type (c6a.48xlarge) with default settings. OrthoFinder computed hierarchical orthologous groups (HOGs) for each internal node in the species tree. To improve HOG prediction accuracy, an outgroup proteome (Burkholderia) was used to root the resulting species tree. HOGs are sets of proteins descended from a single gene in the ancestral species corresponding to the respective internal node. In this study, we focused on analyzing HOGs associated with the species tree node representing the last common ancestor of all A. Pseudomonas, Acinetobacter, and Burkholderia. Specifically, we examined HOGs containing cbrA, crc, and members of the *pel*, *cupB*, and *cupC* operons. For visualizing and annotating phylogenetic trees, custom Python scripts were employed to generate the datasets for annotation in the Interactive Tree of Life (iTol) tool (https://itol.embl.de/). Jupyter notebooks for downloading genomes from NCBI, processing OrthoFinder results, and creating figures are available on GitHub at https://github.com/JonWinkelman/dash_app_pseudomonas.

## ACKNOWLEDGEMENTS

We thank Eric Littman (Duchossois Family Institute) and the Genomics Core Facility at the University of Chicago for help with whole genome sequencing. RNA sequencing was performed by Microbial Genome Sequencing Center (MiGS). Bioinformatics analysis was carried out by Trestle LLC. We thank Jonathan Winkelman for help with generating the phylogenetic tree and Dan Kearns for comments on the manuscript. This work was supported by the NIH Grant R35GM139537 to C.M.W., and NIH Grant R00GM129424 and the Searle Scholars Program Grant SSP-2022-104 to S.M.

## CONFLICT OF INTEREST

The authors declare that they have no conflict of interest.

## SUPPLEMENTAL FIGURES

**Supplemental Fig. 1:**
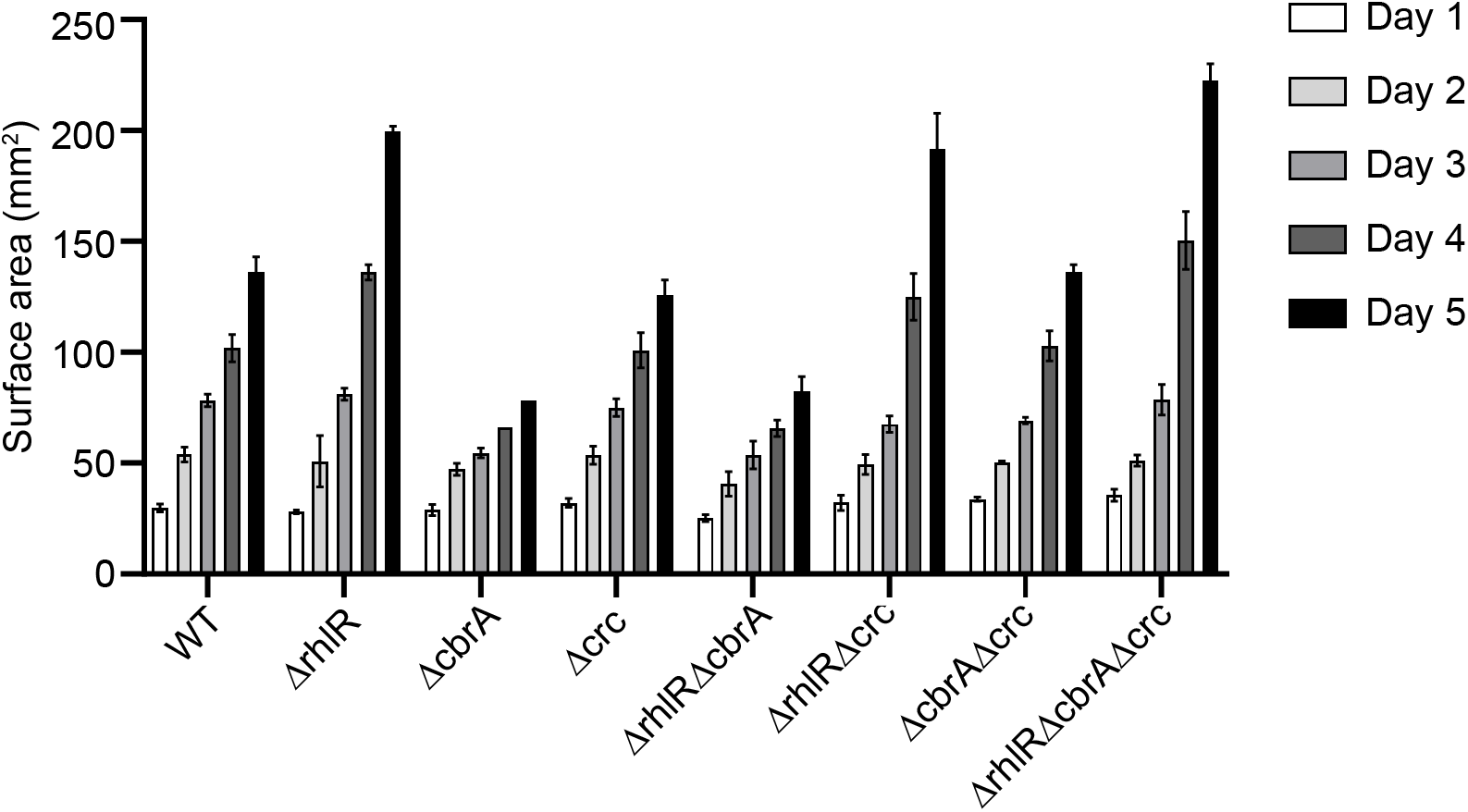
Increase in biofilm surface area coverage of Δ*cbrA* and Δ*rhlR*Δ*cbrA* mutants is slower compared to WT and other studied mutants. Five-day time courses showing development of the colony biofilm surface area of the WT and indicated mutant strains. Error bars represent standard deviation of three biological replicates.

**Supplemental Fig. 2:**
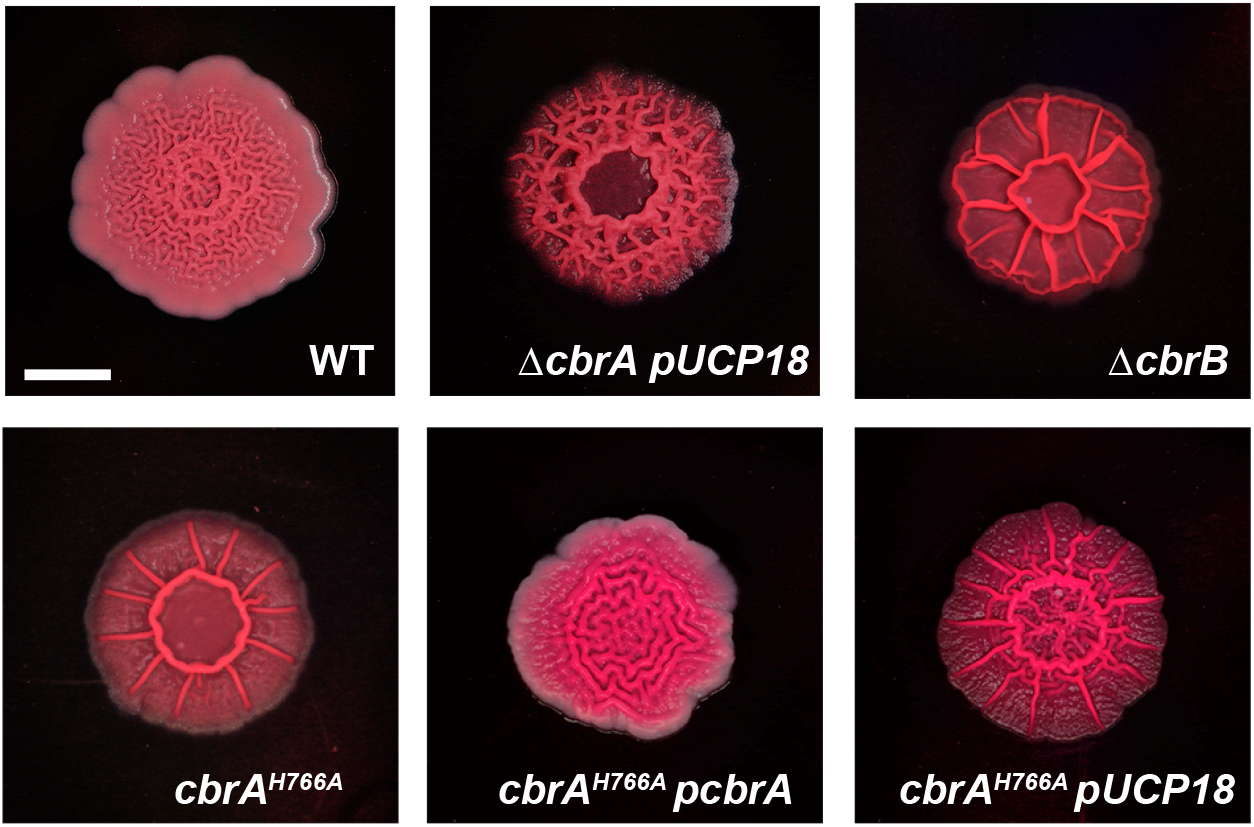
CbrA mediated repression of biofilms requires its cognate response regulator CbrB and its kinase activity. Colony biofilm phenotypes of the designated mutants on Congo red agar medium after 120 h of growth. Scale bar, 5 mm.

**Supplemental Fig. 3:**
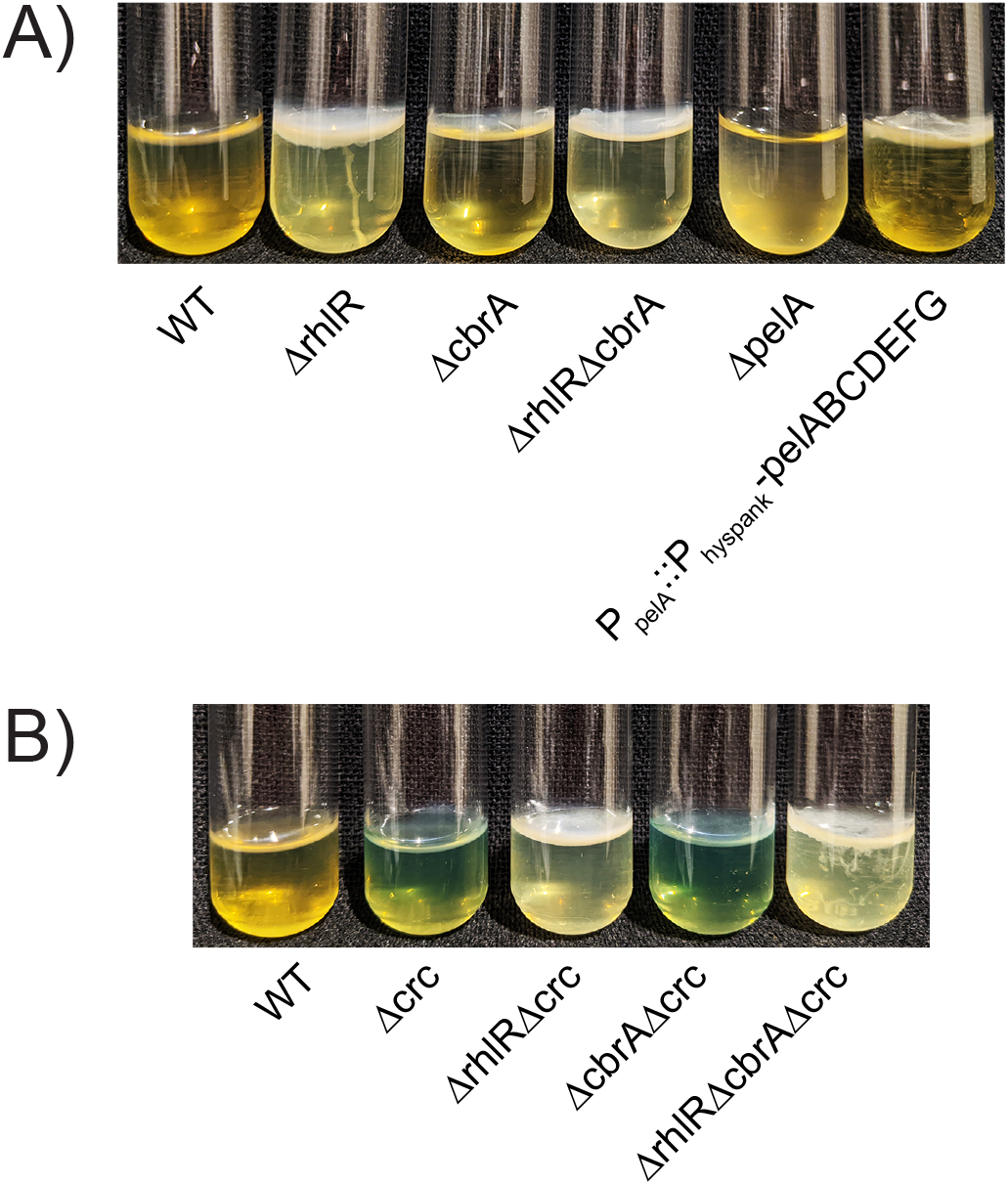
CbrA and RhlR pathways repress pellicle development in addition to colony biofilms. Pellicle biofilms of the indicated *P. aeruginosa* mutants imaged after 72 h of growth in standing cultures.

**Supplemental Fig. 4:**
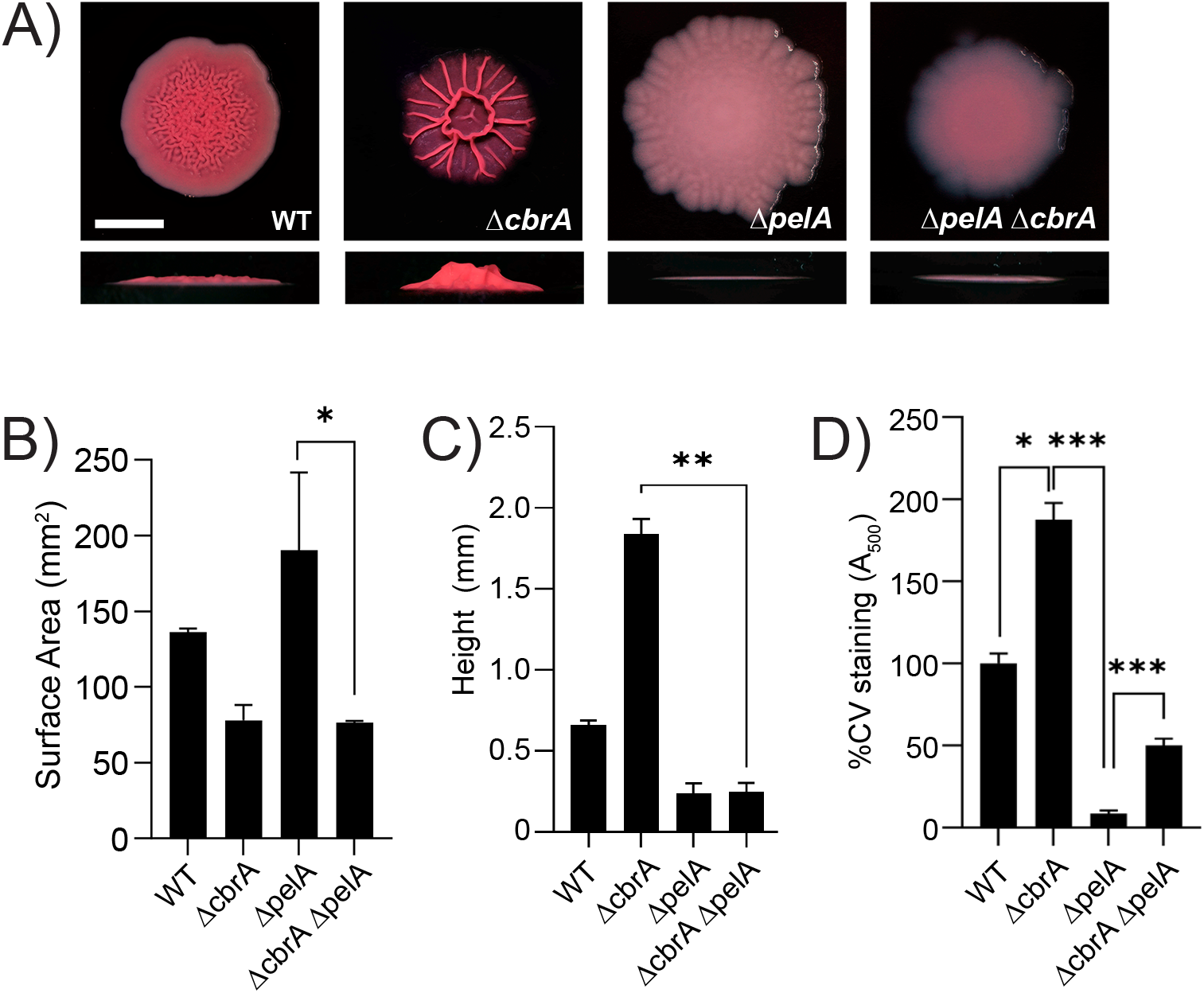
Pel polysaccharide contributes to biofilm formation of the Δ*cbrA* mutant. A) Colony biofilm phenotypes of WT PA14 and the designated mutants on Congo red agar medium after 120 h of growth. Scale bar, 5 mm. B) Colony biofilm surface area quantitation for the indicated strains after 120 h of growth. Error bars represent standard deviation of three independent experiments. Statistical significance was determined using two-tailed *t*-test with Welch’s correction in GraphPad Prism software. * P <0.05. C) Colony biofilm height quantitation for the indicated strains after 120 h of growth. Error bars represent standard deviation of three independent experiments. Statistical significance was determined using two-tailed *t*-test with Welch’s correction in GraphPad Prism software. ** P <0.01. D) Biofilm crystal violet staining assays for WT and indicated mutant strains. Error bars represent standard deviation of three biological replicates. Statistical significance was determined using two-tailed *t*-test with Welch’s correction in GraphPad Prism software. *** P <0.001, * P <0.05.

**Supplemental Fig. 5:**
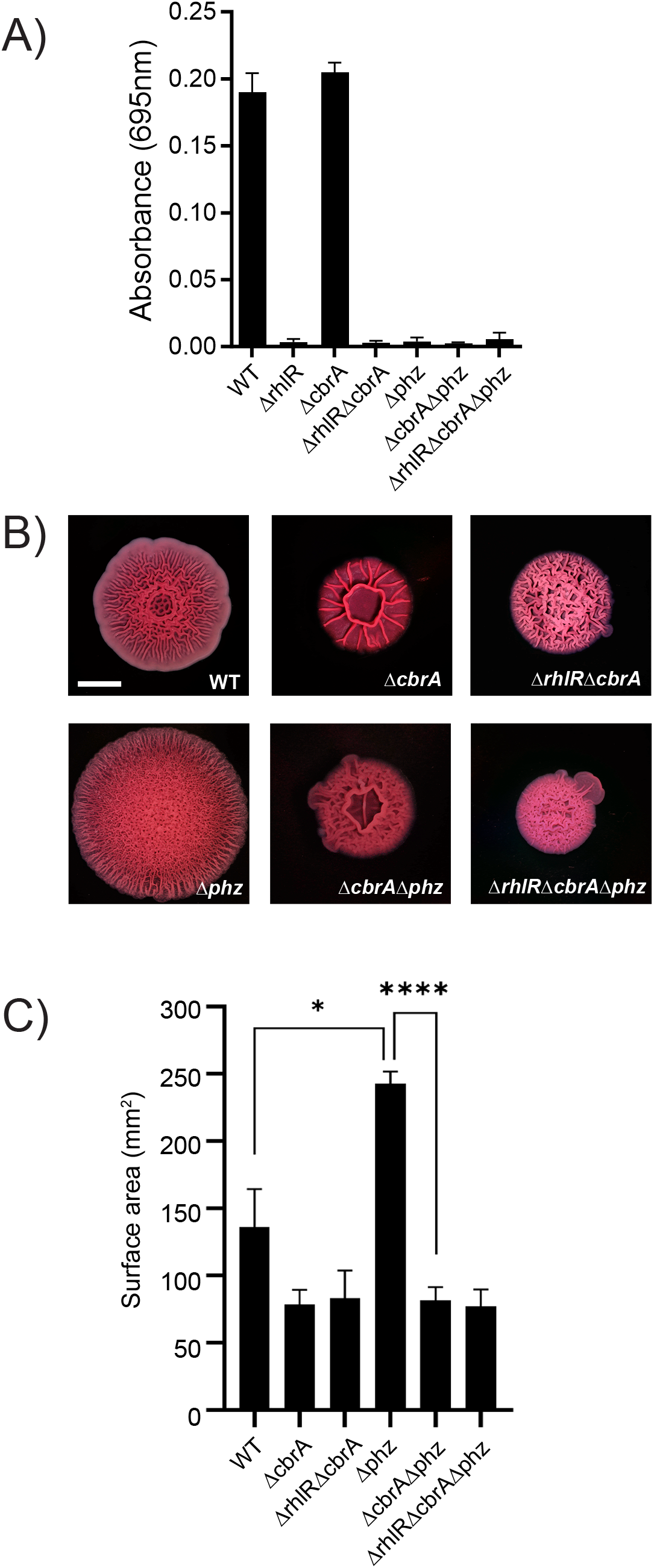
Absence of phenazines is not epistatic to the combined absence of RhlR and CbrA. A) Pyocyanin production phenotypes of the WT PA14 and indicated mutants imaged after overnight growth in LB. B) Colony biofilm phenotypes of the designated mutants on Congo red agar medium after 120 h of growth. Scale bar, 5 mm. C) Colony biofilm surface area quantitation for the indicated strains after 120 h of growth. Error bars represent standard deviation of three independent experiments. Statistical significance was determined using two-tailed *t*-test with Welch’s correction in GraphPad Prism software. ** P <0.01, * P <0.05.

**Supplemental Fig. 6:**
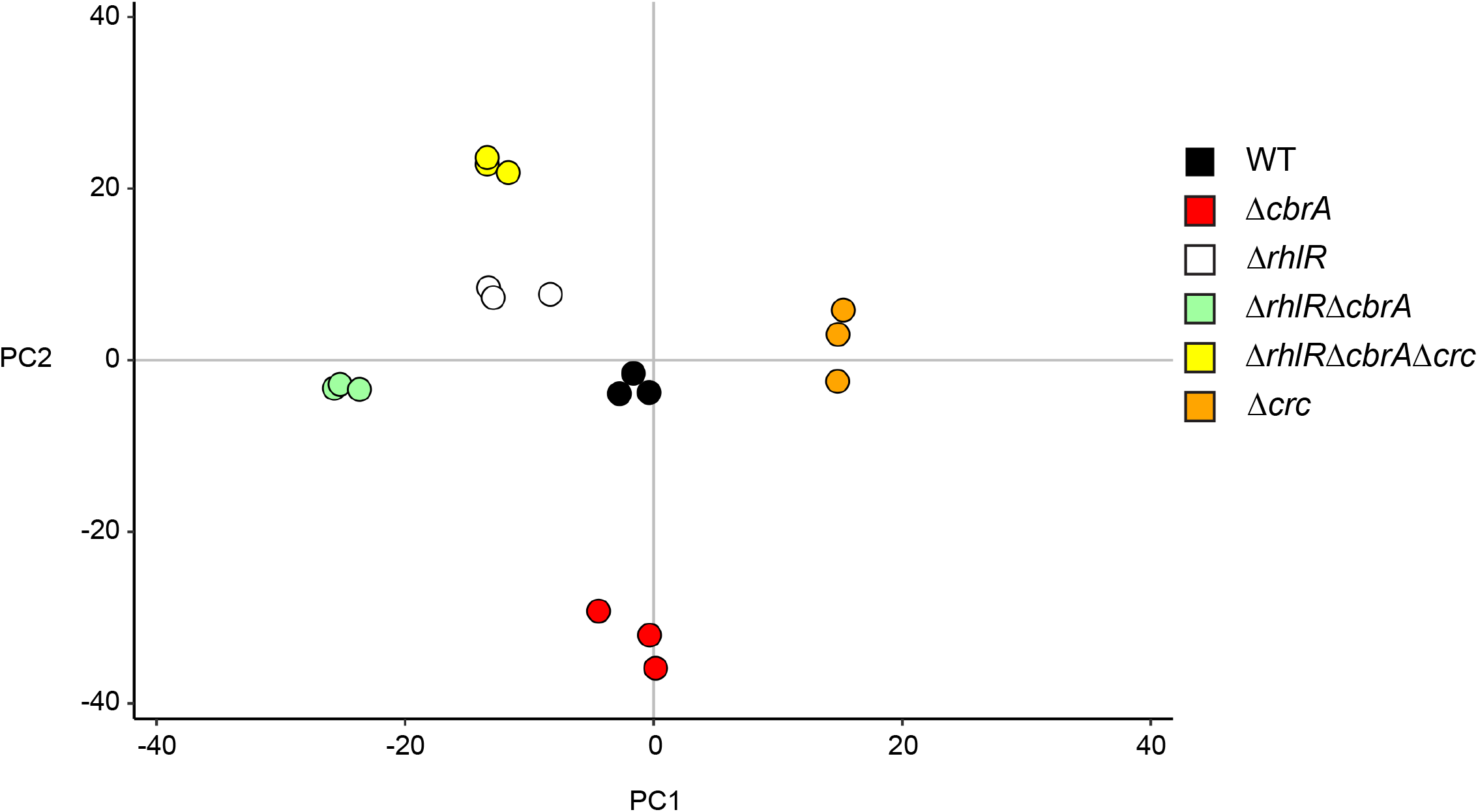
**Principal component analysis (PCA) for RNA-seq on biofilm samples from the indicated strains.**

**Supplemental Fig. 7:**
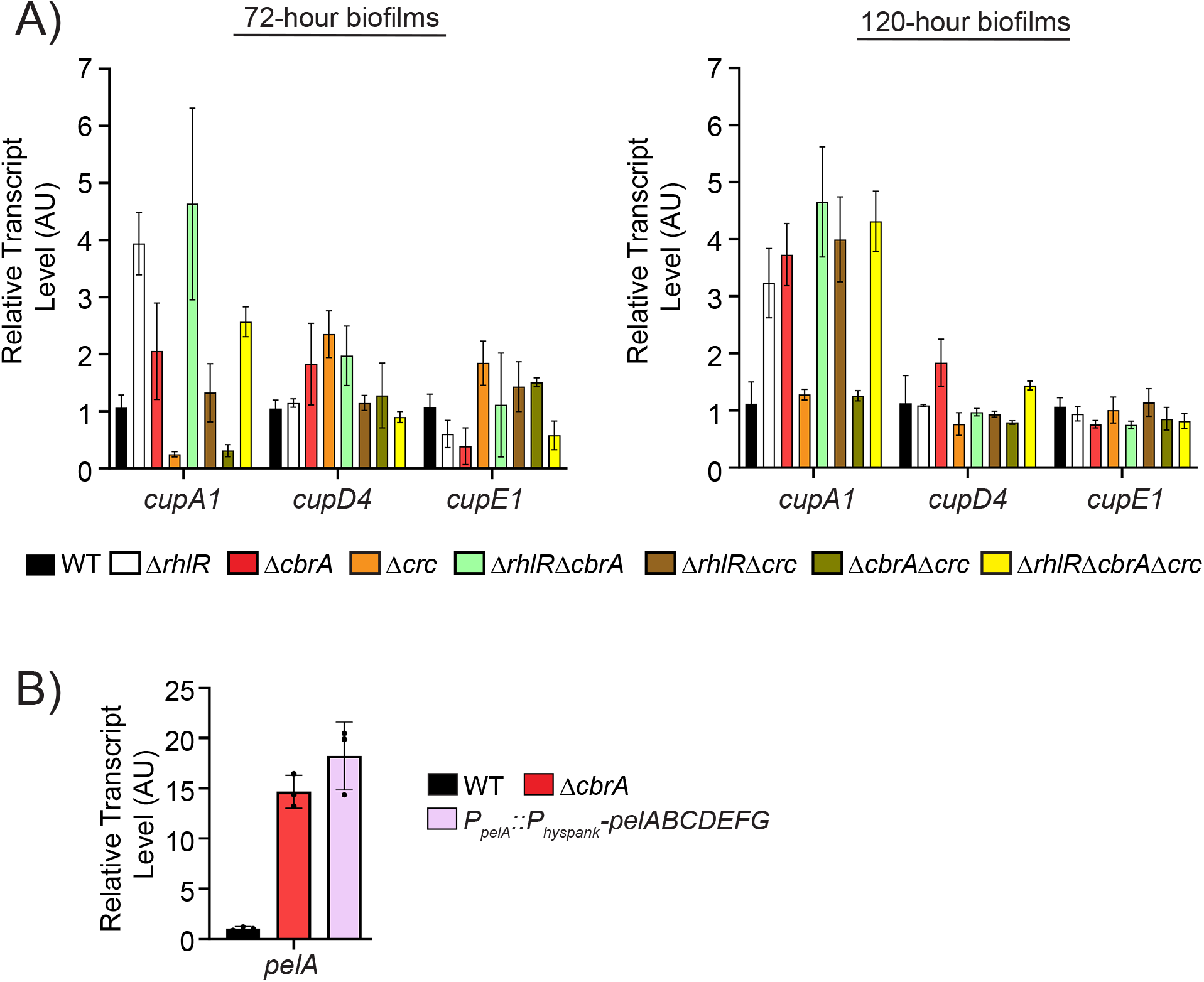
Expression of genes encoding biofilm matrix components in WT and mutant strains in this study. A) Relative expression of *cupA1*, *cupD4*, *cupE1* genes normalized to 16S RNA, *ostA* and *rpsO* transcript levels measured by qRT-PCR in WT PA14 and indicated mutants after 72 h and 120 h of colony biofilm growth. AU denotes arbitrary units. Error bars represent standard deviation of three biological replicates. B) Relative expression of *pelA* gene normalized to 16S RNA, *ostA* and *rpsO* transcript levels measured by qRT-PCR in WT PA14 and indicated mutants after 72 h of colony biofilm growth. AU denotes arbitrary units. Error bars represent standard deviation of three biological replicates.

**Supplemental Fig. 8:**
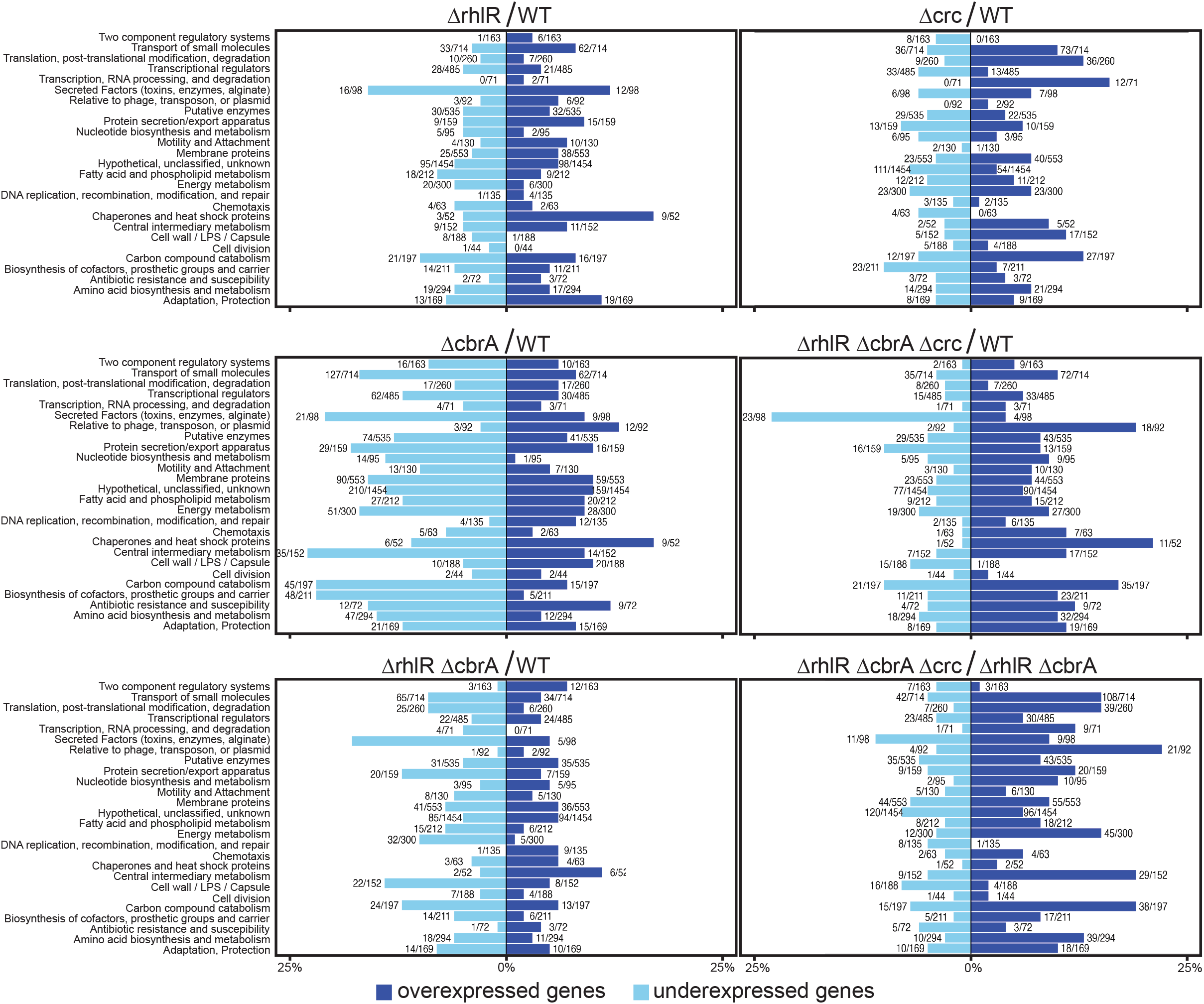
Gene Ontology analysis of RNA-seq data. Percentage of genes in each PseudoCAP category that are downregulated (light blue) or upregulated (dark blue) in Δ*rhlR,* Δ*cbrA,* Δ*crc*, Δ*rhlR*Δ*cbrA* or Δ*rhlR*Δ*cbrA*Δ*crc* compared to WT and Δ*rhlR*Δ*cbrA*Δ*crc* compared to Δ*rhlR*Δ*cbrA*. Numbers next to bars indicated the number of genes up- or downregulated and total number of genes in each PseudoCAP category.

## SUPPLEMENTAL TABLES

**TABLE S1:** Unique suppressor mutations of the Δ*rhlR*Δ*cbrA* biofilm phenotype.

| Strain | PA14 ID<sup>a</sup> | Gene name | Nucleotide position | Mutation |
| --- | --- | --- | --- | --- |
| SPa1068 | PA14_70390 | *crc* | 6275079 | +T (frameshift) |
| SPa1070 | PA14_70390 | *crc* | 6276133 | +T (frameshift) |
| SPa1073 | PA14_70390 | *crc* | 6275042 | M1I (ATG→ATA) |
| SPa1074 | PA14_70390 | *crc* | 6275704 | +G (frameshift) |
| SPa1078 | PA14_70390 | *crc* | 6275207 | +G (frameshift) |
| SPa1079 | PA14_70390 | *crc* | 6275254 | Δ2 bp (frameshift) |
| SPa1088 | PA14_08920 | *rplP* | 763531 | R82P (CGT→CCT) |
| SPa1090 | PA14_08850 | *rplC* | 759596 | T13N (ACC→AAC) |
| SPa1092 | PA14_70390 | *crc* | 6275734 | Δ55 bp (frameshift) |
| SPa1093 | PA14_70390 | *crc* | 6275223 | E61STOP (GAG→TAG) |
| SPa1095 | PA14_70390 | *crc* | 6275107 | W22STOP (TGG→TAG) |
| SPa1096 | PA14_70390 | *crc* | 6275757 | Δ9 bp |

a: annotation from www.pseudomonas.com (Winsor et al., 2016)

**TABLE S2:**
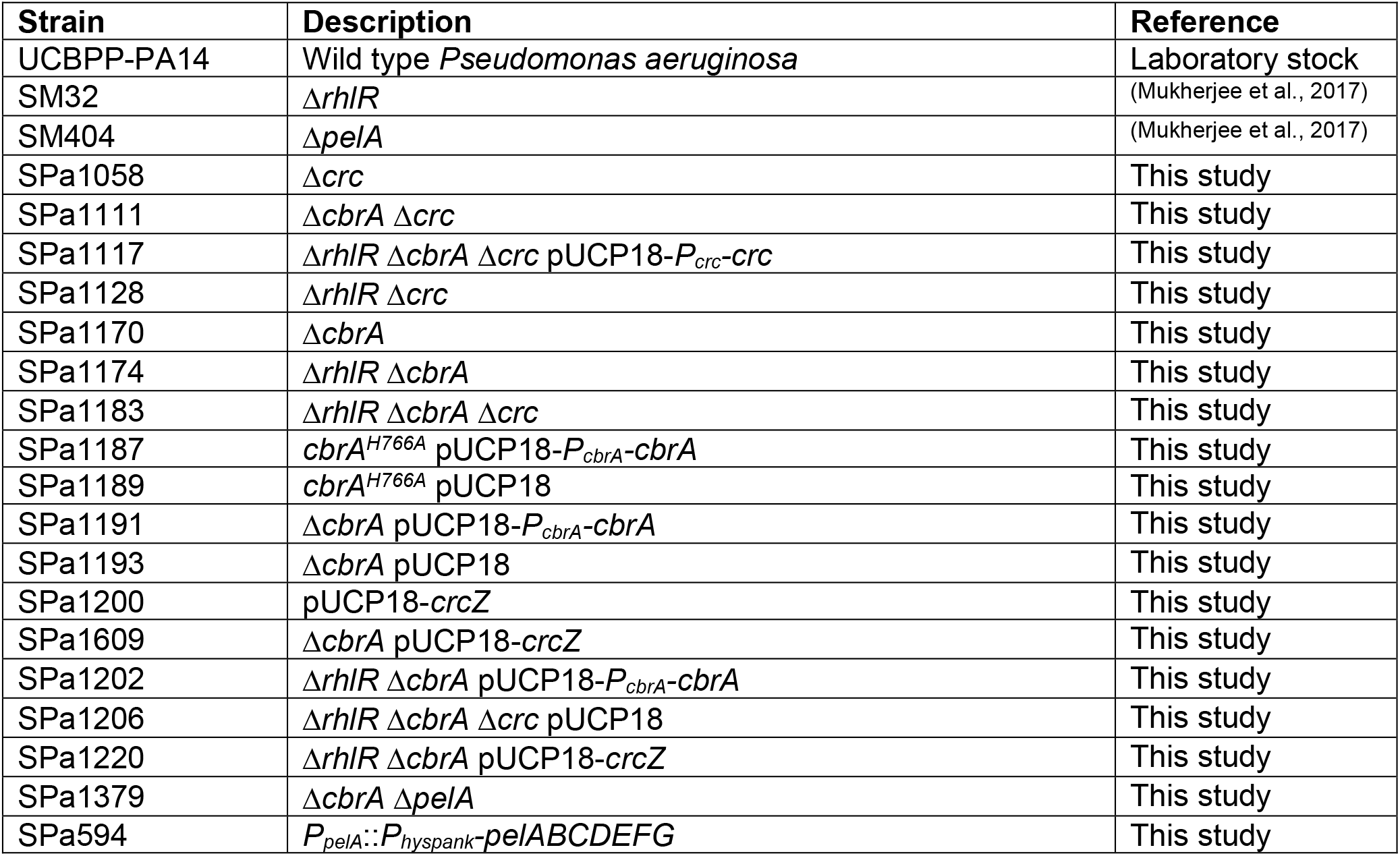
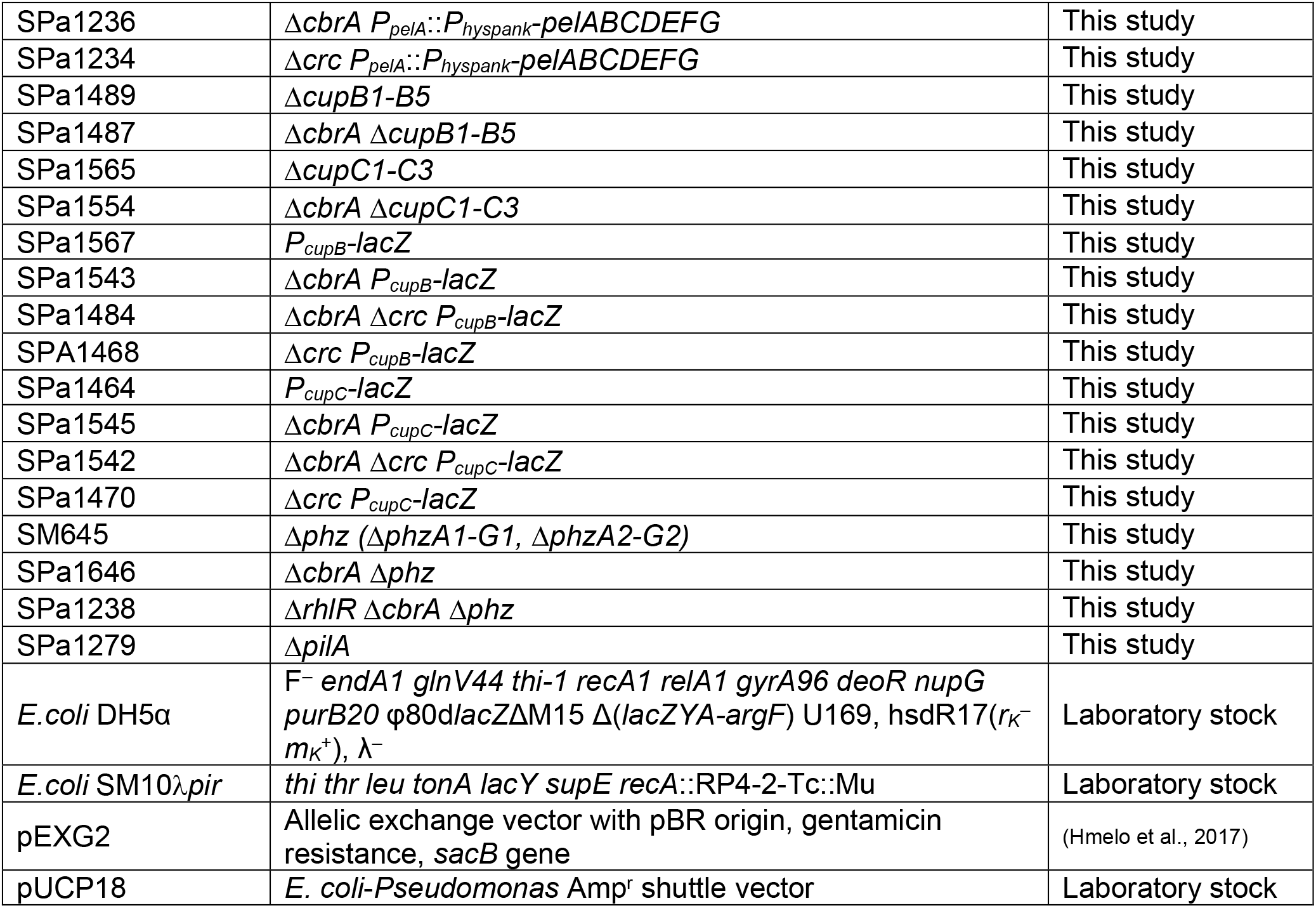
Bacterial strains and plasmids.

**TABLE S3:** RNAseq datasets.

**TABLE S4:** Crc binding site prediction in selected target transcripts.

## Notes

### Competing Interest Statement

The authors have declared no competing interest.

### Summary of Updates

Authors updated; Data, figures and supplemental files updated.

